# BCG administration promotes the long-term protection afforded by a single-dose intranasal adenovirus-based SARS-CoV-2 vaccine

**DOI:** 10.1101/2023.03.21.533720

**Authors:** Dilhan J. Perera, Pilar Domenech, George Giorgi Babuadze, Maedeh Naghibosadat, Fernando Alvarez, Cal Koger-Pease, Lydia Labrie, Matthew Stuible, Yves Durocher, Ciriaco A. Piccirillo, André Lametti, Pierre Olivier Fiset, Seyyed Mehdy Elahi, Gary P. Kobinger, Rénald Gilbert, Martin Olivier, Robert Kozak, Michael B. Reed, Momar Ndao

**Affiliations:** Division of Experimental Medicine, McGill University; Montréal, QC, Canada; Infectious Diseases and Immunity in Global Health Program, Research Institute of the McGill University Health Centre; Montréal, QC, Canada; McGill International TB Centre, McGill University; Montréal, QC, Canada; Department of Biological Sciences, Sunnybrook Research Institute, University of Toronto; Toronto, ON, Canada; Department of Microbiology and Immunology, McGill University; Montréal, QC, Canada; Department of Pathology, McGill University; Montréal, QC, Canada; Department of Production Platforms & Analytics, Human Health Therapeutics Research Center, National Research Council Canada; Montréal, QC, Canada; Département de Microbiologie-Infectiologie et Immunologie, Faculté de Médecine, Université Laval; Québec, QC, Canada; Department of Laboratory Medicine and Molecular Diagnostics, Division of Microbiology, Sunnybrook Health Sciences Centre; Toronto, ON, Canada; National Reference Centre for Parasitology, McGill University Health Centre; Montréal, QC, Canada

**Author notes:** These authors contributed equally (Dilhan J. Perera, Pilar Domenech, George Giorgi Babuadze, Maedeh Naghibosadat, Fernando Alvarez). Corresponding Authors: Dr. Momar Ndao Dr. Michael B. Reed Dr. Robert Kozak.

## Abstract

Despite medical interventions and several approved vaccines, the COVID-19 pandemic is continuing into its third year. Recent publications have explored single-dose intranasal (i.n.) adenovirus-based vaccines as an effective strategy for curbing SARS-CoV-2 in naïve animal models. However, the effects of prior immunizations and infections have yet to be considered within these models. Here, we investigate the immunomodulatory effects of *Mycobacterium bovis* BCG pre-immunization on a subsequent S-protein expressing i.n. Ad vaccination, termed Ad(Spike). We found that Ad(Spike) alone conferred long-term protection from severe SARS-CoV-2 pathology within a mouse model, yet it was unable to limit initial infection 6 months post-vaccination. While i.n. Ad(Spike) retains some protective effect after 6 months, a single administration of BCG-Danish prior to Ad(Spike) vaccination potentiates its ability to control viral replication of the B.1.351 SARS-CoV-2 variant within the respiratory tract. Though BCG-Danish had no effect on the ability of Ad(Spike) to generate and maintain humoral immunity, it promoted the generation of cytotoxic and Th1 responses over suppressive FoxP3^+^ T_REG_ cells in the lungs of infected mice. These data demonstrate a novel vaccination strategy that may prove useful in limiting future viral pandemics by potentiating the long-term efficacy of next generation mucosal vaccines within the context of the safe and widely distributed BCG vaccine.

**One sentence summary:** BCG enhances anti-SARS-CoV-2 immunity and protection afforded by a novel adenovirus-vectored vaccine.

## INTRODUCTION

The COVID-19 pandemic has resulted in over 754 million cases and over 6.8 million deaths as of February 2023^1^. Infection with SARS-CoV-2 can cause a broad spectrum of disease ranging from mild symptoms to severe lung injury and multi-organ failure, potentially leading to death, especially in the elderly and those with comorbidities^2^. Additionally, there is evidence that recovered individuals can experience symptoms termed “long COVID,” which can involve multiple organ systems (e.g., lung, heart, kidneys, liver, etc.)^3^. Consequently, COVID-19 has had severe consequences on global health and the economy and has been the target of novel immunization approaches. Vaccination, in combination with public health measures, has effectively slowed the progression and hospitalization rates associated with SARS-CoV-2^4^; however, these immunization strategies have so far failed to fully prevent viral transmission and infection^5^. Moreover, vaccine efficacy at preventing infection declines by six months after full vaccination^6^, and breakthrough infections, especially by novel SARS-CoV-2 variants, have been reported in previously vaccinated individuals^7^. Indeed, current vaccination efforts have largely focussed on humoral immunity, which wanes over time^8,9^, leading experts to postulate that if long lasting protection is to be achieved, an effective memory T-cell response must be generated against COVID-19 and its variants^10,11^. In addition, there is an urgent need to provide equitable access to affordable and effective vaccines amongst developing nations to prevent the ongoing morbidity and mortality as well as reduce the risk of novel variants emerging.

Among the strategies in preclinical development, a most promising approach involves the use of intranasal (i.n.) vaccines, theoretically capable of eliciting local mucosal immune responses within the respiratory tract that can effectively neutralize SARS-CoV-2 entry and prevent viral replication at the site of initial infection^12^. However, while some vaccines are undergoing phase I/II clinical trials, the success of this approach has thus far been elusive^13^. For example, i.n. administration of a chimpanzee adenovirus-based vaccine carrying the Spike protein (ChAd-SARS-CoV-2-S) successfully generated neutralizing IgA antibodies and T-cell responses in the lungs of hACE-2 mice^14^ and provides at least one month of protection in mice and rhesus macaques^15^. However, a recent progress report from a phase I clinical trial of the i.n. administration of ChadOx1 has announced a failure to substantially increase mucosal immunity in humans^16^. Nonetheless, the safety results were sufficiently encouraging to pursue approaches that enhance mucosal immunity generated using adenovirus vaccine vectors.

In many countries, the live-attenuated *Mycobacterium bovis* Bacillus Calmette-Guérin (BCG) vaccine is administered to newborns soon after birth to protect them from disseminated *Mycobacterium tuberculosis* infection. In fact, most individuals worldwide are BCG vaccinated; of the approximately 140 million babies born per year, approximately 100 million of them are vaccinated with BCG^17^. Curiously, vaccination with BCG has been reported also to reduce child mortality^18^ from neonatal sepsis and lower respiratory tract infections^19^. There is cumulating evidence that BCG acts on the antiviral immune response by boosting the activity of innate immune cells, a concept known as trained immunity^20^, as well as by promoting heterologous T cell activation^21^. As such, many researchers have speculated on the possibility that BCG could provide immunity against SARS-CoV-2 infection^22^, although there are mixed experimental data thus far to support this idea^23,24^. Still, there remains great promise that BCG can be combined with conventional SARS-CoV-2 vaccination strategies to improve their effectiveness^25^. As such, we hypothesized that BCG may provide a novel, cost-effective, and safe priming strategy to enhance the long-term efficacy of i.n. immunization with adenovirus-based COVID-19 vaccines.

Here, we sought to evaluate the effect of prior administration of BCG on the immunogenicity and efficacy of an i.n. human adenovirus serotype 5 (AdV) vaccine expressing the SARS-CoV-2 S (Spike)-protein, referred to as Ad(Spike), in a mouse model of infection with the B.1.351 variant of SARS-CoV-2. Previous work has shown that a single i.n. immunization with adenovirus-based vaccines against SARS-CoV-2 is sufficient to provide effective protection from infection in naïve animals^26^. In this report, we demonstrate that the protection conferred by our intranasally administered Ad(Spike) vaccine alone declines in mice by 6 months post-immunization. Importantly, a single dose of BCG administered prior to Ad(Spike) vaccination was capable of specifically boosting the Spike-specific cytotoxic T-cell response in the lungs and, in doing so, significantly decreasing the replication and production of infectious viral particles up to 6 months after i.n. immunization with the Ad(Spike) vaccine. These results show a novel and innovative approach to the use of heterologous vaccine strategies incorporating the widely approved BCG vaccine to protect against current and future viral pandemics.

## RESULTS

### The *in vivo* protection provided by a single dose of intranasal recombinant Ad(Spike) attenuates with time

To investigate the long-term protection provided by i.n. vaccine administration, we developed a replication-deficient (ΔE1-, ΔE3-) human adenovirus serotype 5 expressing the full-length S-protein of the ancestral strain of SARS-Co-V-2 (Ad(Spike)) which was codon optimized for expression in human and mouse cell lines (Figure 1a). Exogenous gene expression was verified *in vitro* by western blot using S-protein specific antibodies (Figure 1b). Female C57BL/6 mice were vaccinated with either PBS or 10^9^ TCID_50_ Ad(Spike) intranasally. The infectious dose of Ad(Spike) was determined using a pilot study and was chosen based on the induction of Spike-specific IgA antibodies in lung homogenates from vaccinated animals (Supplemental Figure 1). Study animals were challenged 2 or 6 months later with the Beta variant (B.1.351) of SARS-CoV-2. In each case, the animals were followed for 5 days post-infection (dpi) to determine viral shedding from oral swabs (3-dpi) and the viral load in the lungs at 5-dpi (Figure 1c).

**Figure 1.**
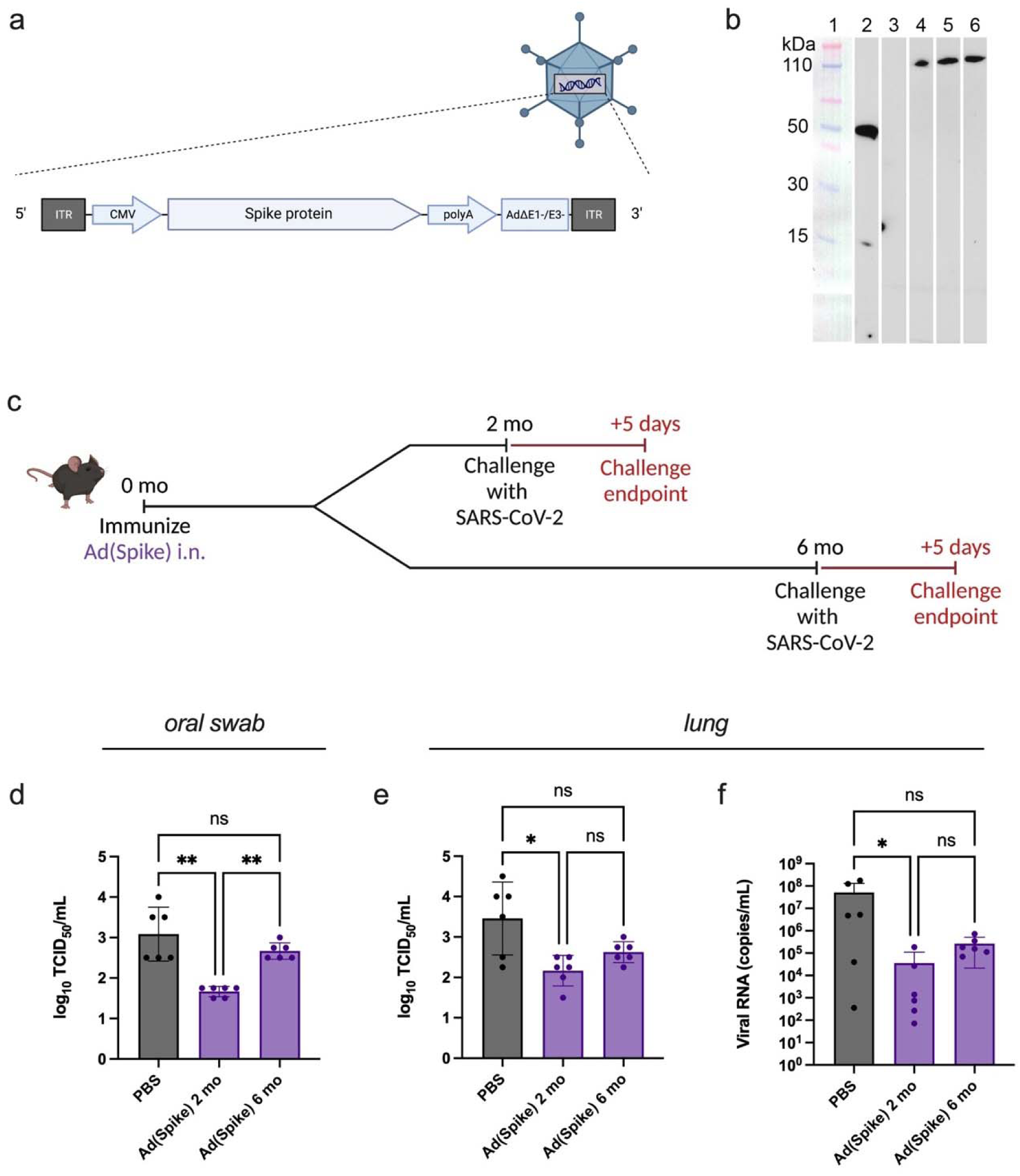
The protection provided by a single dose of intranasal recombinant Ad(Spike) attenuates with time in C57BL/6 mice. (**a**) Diagram of the SARS-CoV-2 Spike protein (ancestral strain) transgene cassette expressed in our recombinant ΔE1/E3 human adenovirus serotype 5, termed Ad(Spike). (**b**) Western blot analyses of antigen expression from SF-BMAd-R cells infected with Ad(Spike). Samples were run against a molecular weight ladder; lanes 1 and 2. The negative control contains SF-BMAd-R cells infected with a ΔE1/E3 adenovirus with an empty gene cassette, termed Ad(e); lane 3. Ad(Spike) was amplified in 3 batches of cells and western blots were run on cell lysates from each batch; lanes 4-6, before being combined and purified. This western blot was run with an RBD-specific antibody, thereby capturing the S1 portion. (**c**) Study design schematic. At time 0, animals were vaccinated intranasally (i.n.) with 10^9^ mean tissue culture infectious dose (TCID_50_) of Ad(Spike) in 30 uL. In the case of the sham control, at time 0 animals were vaccinated i.n. with 30 uL PBS. Mice were then challenged with 10^6^ TCID_50_ SARS-CoV-2 South African strain (B.1.351) at month 2 or month 6 post-vaccination. In both challenge models, animals were followed for 5 days with nasal swab collection on day 3 and euthanasia on day 5. (**d**)-(**f**) Infectious viral load in mice challenged with the B.1.351 variant of SARS-CoV-2, 2- and 6- months post-immunization with Ad(Spike). Viral load (TCID_50_) in (d) oral swabs at 3 dpi, and (e) lungs at 5 dpi, quantified by the Spearman–Kärber method. Viral RNA in mouse (f) lungs at 5 dpi. N=6. Data points represent individual mice, means ± SD are shown. For (d)-(f), Kruskal-Wallis test with Dunn’s multiple comparisons: *p<0.05; **p<0.01; ns = not significant. Schematics made with BioRender.com.

As expected, Ad(Spike)-immunized mice displayed a significant reduction in the production of infectious SARS-CoV-2 in oral swabs 2 months after vaccination (Figure 1d). However, when challenged 6 months post-vaccination, viral titres in vaccinated animals were comparable to unvaccinated mice (Figure 1d). Similarly, infectious viral titres (Figure 1e) and SARS-CoV-2 RNA (Figure 1f) assessed directly in the lungs of infected animals at 6 months post-vaccination were not statistically different from unvaccinated mice as opposed to data from mice challenged 2 months after immunization, where significant differences were observed. These findings reveal that the protection from infection conferred by a single-dose Ad(Spike) i.n. vaccine against SARS-CoV-2 is short-lived, waning over time.

### BCG administration prolongs the protective effect of Ad(Spike) in immunized mice

We next investigated the effectiveness of a prime-boost vaccination regimen using BCG. Female C57BL/6 mice were pre-immunized with 10^6^ colony forming units (CFU) of the BCG-Danish strain containing an empty plasmid (BCG(e)) intraperitoneally (i.p.), 1 month prior to i.n. Ad(Spike) vaccination (month 0). Naïve controls received the vehicle (PBS) in place of both BCG(e) and Ad(Spike). First, we tested the possibility that BCG alone could provide non-specific protection against SARS-CoV-2 in our animal model, since reports from human data are controversial^27^. Animals were pre-immunized with BCG(e) and vaccinated with an adenoviral vector containing an empty gene cassette (Ad(e)) before challenge with SARS-CoV-2 two months later. Mice pre-immunized with BCG-Danish displayed no significant reduction in infectious viral titres or viral RNA in oral swabs or lungs (Supplemental Figure 2) compared to PBS controls, confirming the BCG-Danish vaccine did not provide significant non-specific protection to challenged mice^23,28^.

We then investigated the effect of prior BCG-Danish administration on the duration of our Ad(Spike) vaccine 6 months after vaccination (Figure 2a) by quantifying the daily variation in viral replication and infectious particles in oral swabs and the pulmonary tissue of mice infected with SARS-CoV-2. Here, we found that while a single dose of i.n. Ad(Spike) could reduce infectious viral particles in oral swabs at 1- and 3-dpi, mean viral titres in animals that were first pre-immunized with BCG were significantly lower than those that were not (Figure 2b). At 5 dpi, we assessed also the reduction of viral RNA in oral swabs (Figure 2c), infectious virus in the lungs measured by TCID_50_ (Figure 2d), and viral RNA in the lungs (Figure 2e). We found that by 6 months post-vaccination, although a single dose of Ad(Spike) was able to reduce viral burden, compared to controls, this reduction was not statistically significant. However, significance was rescued in animals that first received BCG(e), revealing that while i.n. Ad(Spike) retains some protective effect after 6 months, a single administration of BCG-Danish prior to Ad(Spike) vaccination potentiates its ability to control viral replication within the respiratory tract.

**Figure 2.**
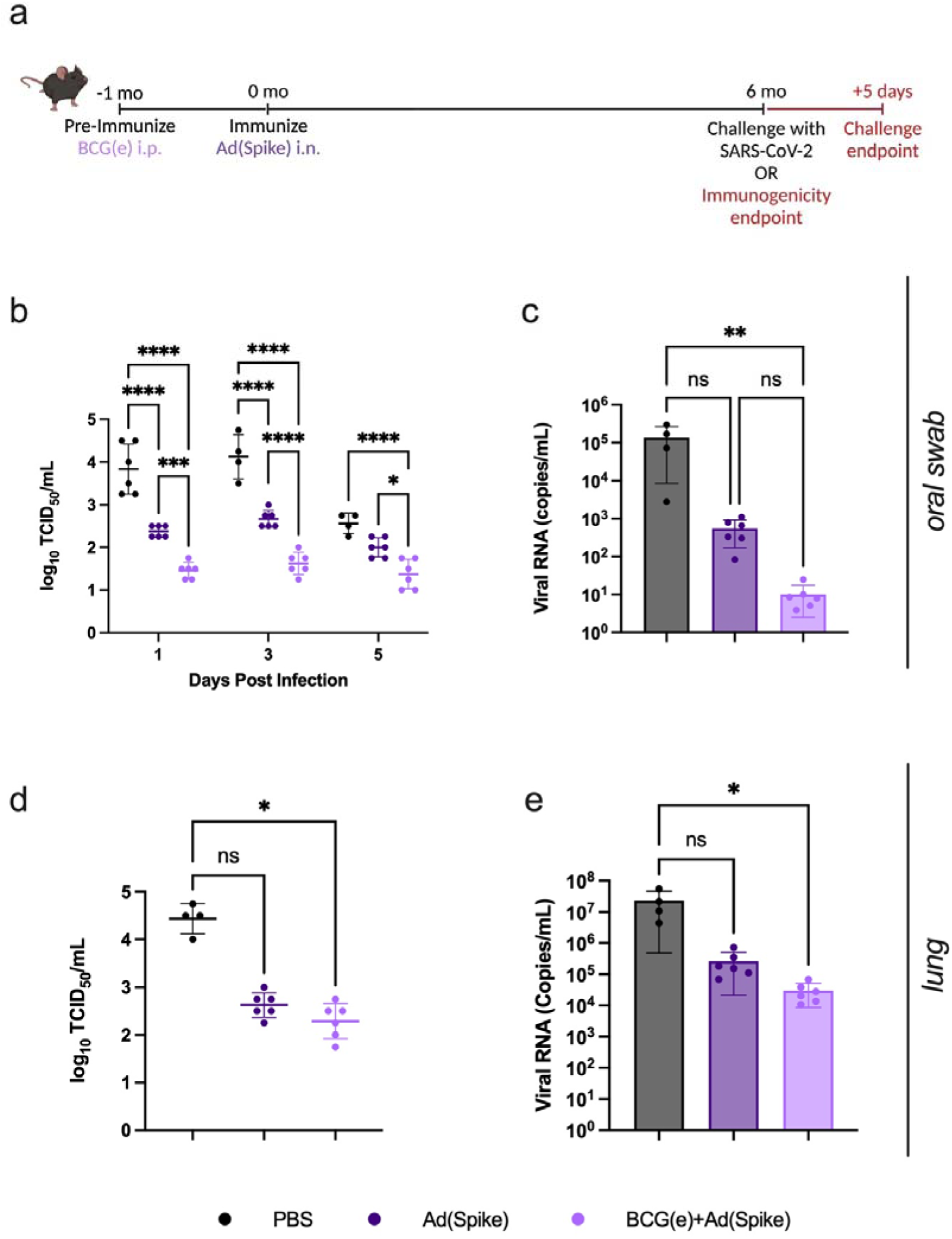
Exposure to BCG prolongs the protective effect of Ad(Spike) in immunized C57BL/6 mice. **(a)** Study design schematic. Animals were primed intraperitoneally (i.p.) with 10^6^ cfu BCG containing an empty gene cassette (BCG(e)) or PBS at time -1 month. At time 0, animals were vaccinated intranasally (i.n.) with 10^9^ mean tissue culture infectious dose (TCID_50_) of Ad(Spike) in 30 uL. In the case of the sham control, at time 0 animals were vaccinated i.n. with 30 uL PBS. Mice were then challenged with 10^6^ TCID_50_ SARS-CoV-2 South African strain (B.1.351) 6 months post-vaccination. Animals were followed for 5 days post challenge, with nasal swabs collected on days 1, 3, and 5 post-infection and euthanasia on day 5. **(b)-(e)** Viral load quantified (b) in oral swabs at 1, 3, and 5 dpi, and (d) in lungs at 5 dpi, quantified by TCID_50_. Viral RNA in (c) oral swabs at 5 dpi and (e) lungs at 5 dpi. N=4-6. Data points represent individual mice, means ± SD are shown. For (b), Two-way ANOVA with Tukey’s multiple comparisons: *p<0.05; ***p<0.001; ****p<0.0001. For (c)-(e), Kruskal-Wallis test with Dunn’s multiple comparisons: *p<0.05; **p<0.01; ns = not significant. Schematic made with BioRender.com.

### Ad(Spike)-vaccinated animals are protected from SARS-CoV-2-induced pathology 6 months post-vaccination

SARS-CoV-2 infection in C57BL/6 mice causes severe pulmonary inflammation characterized by immune cell infiltration, lung atelectasis, and bronchial constriction. Since we observed a greater reduction of viral particle load in BCG-Danish exposed mice, we next assessed the extent of pulmonary damage at 6 months post-Ad(Spike) vaccination, both with and without BCG pre-immunization. Scoring of lung pathology showed an overall significant reduction in cellular and tissue damage (CTL), circulatory/vascular damage (CVL), and inflammatory patterns (RIP) in all Ad(Spike) vaccinated animals that was not observably enhanced through pre-immunization with BCG(e) (Figures 3a-b-c; Supplemental Table 1). However, we could not distinguish the protective effect of BCG-Danish, as both groups displayed a significant reduction in lung pathology. Collectively, these results confirm that while a single i.n. dose of Ad(Spike) vaccine fails to prevent infection, it potently protects against SARS-CoV-2-induced severe lung pathology as long as 6 months post-vaccination.

**Figure 3.**
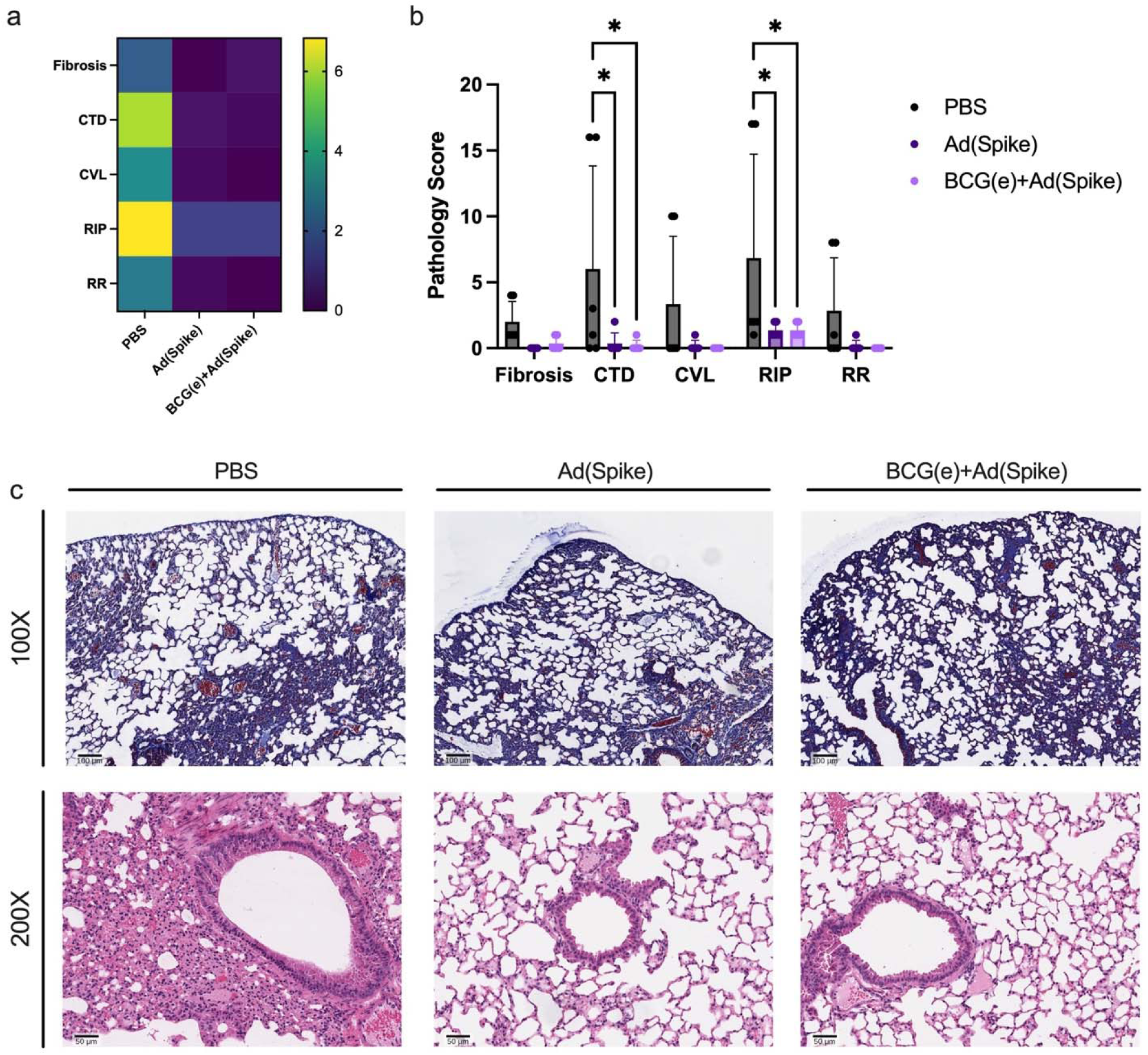
SARS-CoV-2 induced pulmonary pathology is well prevented in Ad(Spike) immunized mice, regardless of BCG exposure. **(a)** Heat map and **(b)** graphical summary of lung pathology (5 days post infection (dpi)) scored in categories: Fibrosis, CTD, CVL, RIP, RR. CTD: Cell/tissue damage which is comprised of bronchoepithelial necrosis (scored 1–3), inflammatory cells/debris in bronchi (1–3), intraepithelial neutrophils (1–3) alveolar emphysema (Yes=1/No=0). CVL: Circulatory/vascular lesions comprised of alveolar hemorrhage (Y/N), significant alveolar edema (Y/N), endothelial/vasculitis (1–3). RIP: Reaction/inflammatory patterns comprised of necrosis/suppurative bronchitis (Y/N), intra-alveolar macrophages (Y/N), mononuclear inflammation around airways (Y/N), neutrophilic/heterophilic inflammation (1–3), mesothelial reaction (1–3). RR: Regeneration/repair, includes alveolar epithelial cell regeneration/proliferation (1–3) and bronchiolar epithelial cells regeneration/proliferation (1–3). **(c)** Lungs of vaccinated animals, infected after 6 months, were harvested at 5 dpi and stained with Masson’s trichrome for fibrosis (top row) and H&E staining for pathology scoring (bottom row). Row one is imaged at 100X magnification (scale bar, 100 µm). Row two is imaged at 200X magnification (scale bar, 50 µm) showing airway mononuclear inflammation in control animals. Each image is representative for the group. N=4-6. Data points represent individual mice, means ± SD are shown. For (b), Two-way ANOVA with Tukey’s multiple comparisons: *p<0.05.

### Ad(Spike)-induced antibody profiles are not influenced by pre-immunization with BCG

Vaccine protection against SARS-CoV-2 infection is largely attributed to its ability to generate high quantities of neutralizing antibodies, although the potential role of BCG in promoting antibody production remains to be assessed. To determine if BCG influenced the quantity and affinity of the antibody response generated by Ad(Spike), we bled immunized animals at select timepoints pre- and post-vaccination. Antibody titres against the full Spike protein (Figure 4) and the receptor binding domain (RBD) (Supplemental Figure 3) were assessed via ELISA. Expectedly, animals in the PBS and BCG(e)+Ad(e) groups did not produce Spike or RBD-specific serum antibodies throughout the study. Spike-specific IgG was detectable at high levels 1 month after vaccination and continued to rise until 2 months after vaccination when levels slowly declined until the end of the study for both Ad(Spike) and BCG(e)+Ad(Spike) groups (Figure 4a). Although RBD-specific IgG was significantly higher in Ad(Spike) animals than those that received BCG(e)+Ad(Spike) at 1 month post-vaccination, this difference was not present at later time points (Supplemental Figure 3a). As such, Spike-specific IgG was similar between vaccinated groups.

**Figure 4.**
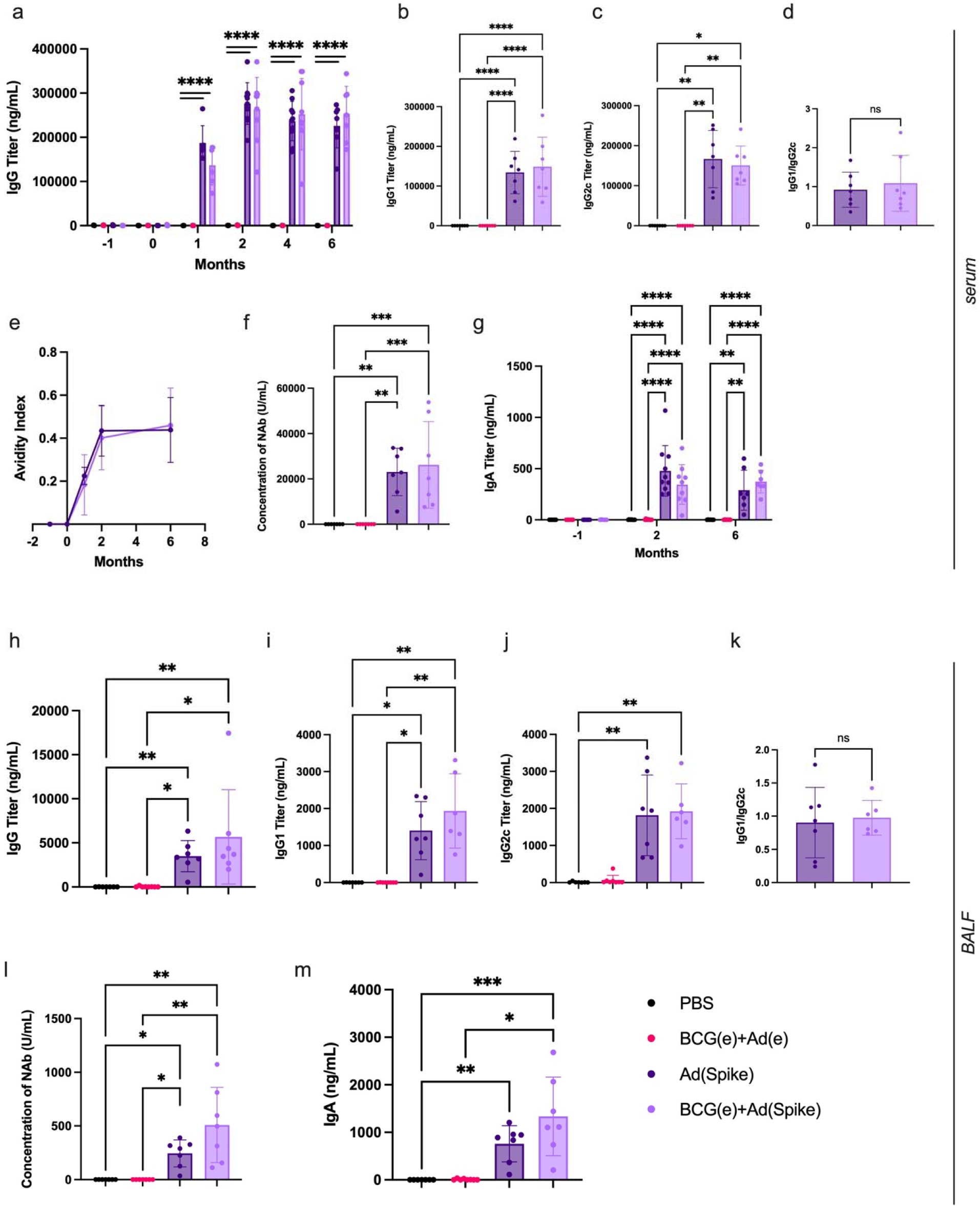
Exposure to BCG does not significantly influence the quantity or quality of Ad(Spike) generated antibodies. Spike-specific antibodies in the (a)-(g) serum and (h)-(m) bronchoalveolar lavage fluid (BALF). (**a**) IgG titers in mouse sera throughout the study schedule determined by ELISA. (**b**) IgG1 and (**c**) IgG2c at 6 months post-vaccination. The ratio of Spike-specific IgG1/IgG2c at 6 months post vaccination is given in (**d**). (**e**) IgG avidity index at -1, 0-, 1-, 2-, and 6-months post vaccination. (**f**) cPass determined antibody neutralization activity in serum at 6 months post vaccination. (**g**) IgA titers in mouse sera calculated at 0-, 3-, and 6-months post vaccination. N=7-10. Spike-specific (**h**) IgG, (**i**) IgG1, and (**j**) IgG2c with the ratio of IgG1/IgG2c given in (**k**). (**l**) cPass determined antibody neutralization activity in BALF at 6 months post vaccination. (**m**) IgA in BALF at 6 months post vaccination calculated by ELISA. N=7. Data points represent individual mice, means ± SD are shown. The included legend applies to both serum and BALF data. For (a), (g), Two-way ANOVA with Tukey’s multiple comparisons: **p<0.01; ****p<0.0001. For (b), (f), (k), One-way ANOVA with Tukey’s multiple comparisons: **p<0.01; ***p<0.001; ****p<0.0001; ns = not significant. For (c), (d), (h)-(j), (l), (m), Kruskal-Wallis test with Dunn’s multiple comparisons: *p<0.05; **p<0.01; ***p<0.001.

Since BCG is known to induce IFNγ^29^, a known promoter of IgG2c production^30^, we then addressed if BCG influenced the isotype of antibodies produced upon Ad(Spike) vaccination in the animals prior to challenge. Both groups of vaccinated animals similarly expressed Spike-specific IgG1 (Figure 4b) and IgG2c (Figure 4c) and displayed the same ratio of IgG1/IgG2c (Figure 4d). Correspondingly, both groups of vaccinated animals expressed similar levels of RBD-specific IgG1 (Figure S3d) and IgG2c (Figure S3e) in similar ratios (Supplemental Figure 3f), confirming that BCG-Danish pre-immunization did not influence isotype-switching upon Ad(Spike) vaccination.

We then assessed IgG avidity by ELISA. As expected, Spike- and RBD-specific IgG avidity rose from 0- to 2-months (Figure 4e and Supplemental Figure 3b, respectively) and Spike-specific avidity was maintained in both vaccinated groups until 6 months, demonstrating that BCG did not influence the avidity of IgG. Interestingly, RBD-specific IgG avidity dropped at 6-months to levels similar to those observed at one-month post-vaccination (Supplemental Figure 3c), suggesting that neutralisation of the ACE2-Spike binding domain declines with time in both groups. In addition, a cPass surrogate virus neutralization assay confirmed that BCG did not influence the abundance of neutralizing antibodies within immunized mice (Figure 4f) compared to mice vaccinated with Ad(Spike) alone. Intranasal vaccination with Ad(Spike) also resulted in detectable levels of antigen-specific serum IgA in both groups (Figure 4g), prompting us to investigate levels of mucosal antibodies. Bronchoalveolar lavage fluid (BALF)-derived Spike-specific IgG antibodies were present and statistically greater than negative controls at 2 and 6 months post-vaccination (Figure 4h). This was observed also for RBD-specific IgG responses in BALF prior to infection (Figure S3h). Specifically, Spike-specific IgG1 (Figure 4i) and IgG2c (Figure 4j), as well as their ratio (Figure 4k), did not differ between the two vaccinated groups. The same pattern was observed for RBD-specific IgG1 (Figure S3i), IgG2c (Figure S3j), and the ratio of the two (Figure S3k). The mean neutralizing activity of BCG-pre-immunized and vaccinated animals was two-fold greater than those animals that were solely Ad(Spike) vaccinated, though this difference was not significant (Figure 4l). This trend of higher antibody levels in the BALF from pre-immunized and vaccinated animals was again seen with the Spike-specific IgA (Figure 4m) and RBD-specific IgA (Figure S3l) titres; though, again the difference was not statistically significant. Taken together, these data show that pre-immunization with BCG-Danish does not promote the persistence of Ad(Spike)-induced protection by modulating the generation, quantity, or quality of circulating or mucosal humoral responses.

### BCG pre-immunization promotes long-term cellular immunity in the lungs

An important aspect of vaccination is to generate robust and lasting tissue-resident memory T cells (T_RM_) in order to confer protection^31^. Since we observed that BCG-Danish did not impact the long-term protective antibody response Ad(Spike) generated against SARS-CoV- 2, we next investigated if BCG-Danish pre-immunization potentiated the generation of memory CD4^+^ and CD8^+^ T_RM_ cells in infected mice. Six months post-vaccination, isolated lung T cells were activated with SARS-CoV-2 Spike protein peptides (Figures 5a; S4). As shown in Figure 5b, at 6 months post-vaccination, the frequencies of lung CD8^+^ T cells from immunized groups were significantly greater than controls regardless of BCG administration. However, when we assessed the production of Granzyme B (GrB) and IFNγ in activated CD69^+^ CD8^+^ T_RM_ cells, we observed that mice that received BCG(e) prior to immunization with Ad(Spike) produced a higher frequency of GrB- (Figure 5c) and IFNγ-secreting CD8^+^ T cells (Figure 5d) compared to the singly vaccinated group 6 months post-vaccination, suggesting that their responses are more cytotoxic in nature. In parallel, we observed an increase in activation (CD69^+^) of CD4^+^ T cells upon peptide restimulation in the immunized group that was previously exposed to BCG (Figure 5e). In fact, mice pre-immunized with BCG, regardless of Ad(Spike) vaccination, had increased frequencies of IFNγ^+^ CD4^+^ (Figure 5f) and IL17A^+^ CD4^+^ T cells in the lung (Figure 5g), although these trends did not reach statistical significance. Coincidently, we observed also significantly less FoxP3^+^ regulatory T (T_REG_) cells among CD69^+^ CD4^+^ T cells (Figure 5h), suggesting that BCG promotes the generation of cytotoxic over suppressive T-cell responses in BCG(e)+Ad(Spike)-immunized mice relative to the other treated groups. Collectively, these results demonstrate that BCG pre-immunization acts on the long-term potency of the Ad(Spike) vaccine by promoting the generation of cytotoxic and Th1 responses over suppressive FoxP3^+^ T_REG_ cells in the lungs of infected mice.

**Figure 5.**
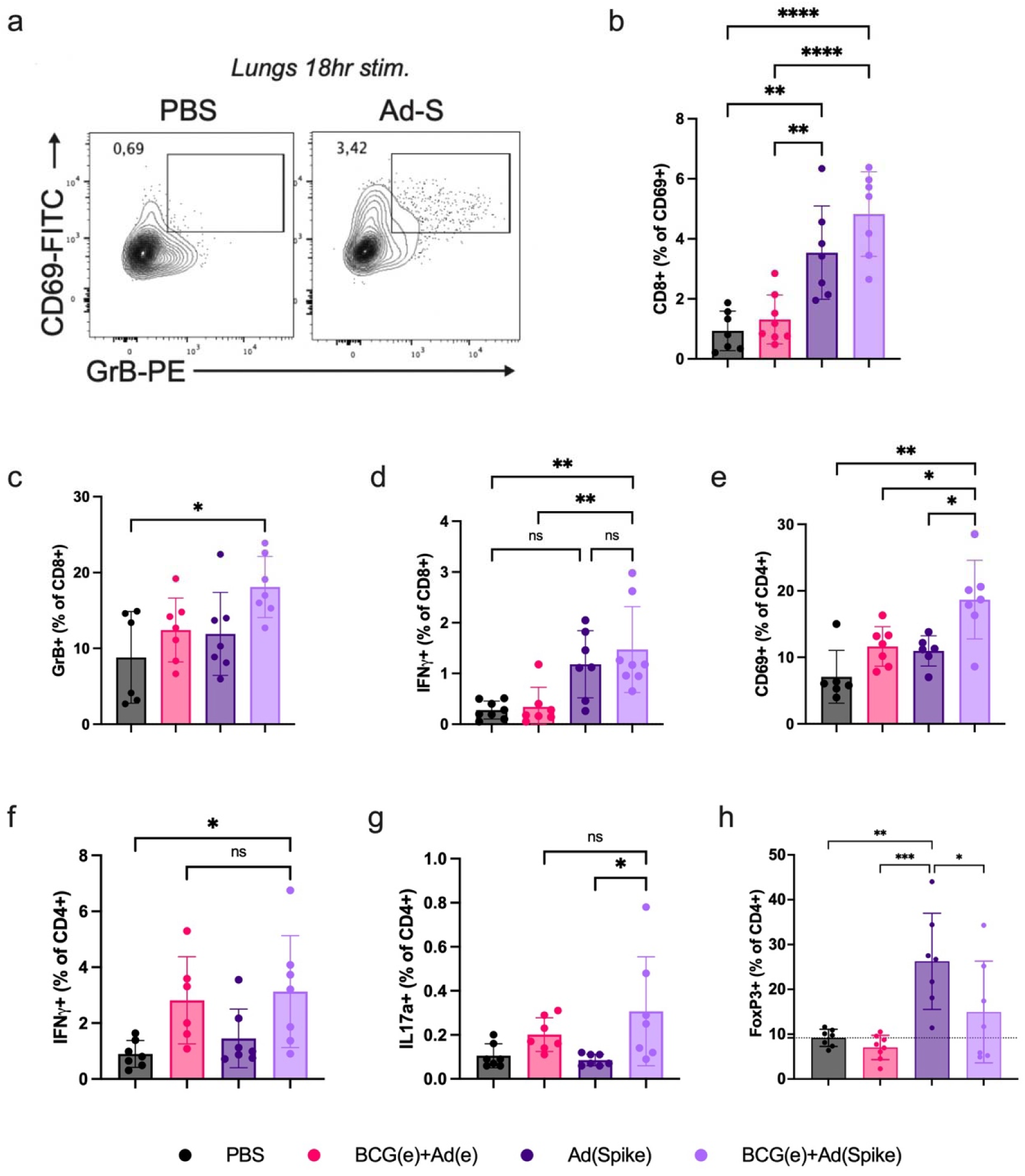
BCG potentiates the generation of long-lasting cellular immunity. Six months post-vaccination, cell mediated responses from lung cells were analysed following 18 hours restimulation with an S-protein peptide pool. (**a**) Expression of CD69 and GrB in vaccinated animals compared to controls (Ad-S=Ad(Spike)). (**b**) Total frequency of CD69+CD8+ T cells. The frequency of CD69+CD8+ T cells which are (**c**) GrB+ and (**d**) IFN□+. (**e**) Frequency of CD69+ cells among the CD4+ population. (**f**) IFN□, (**g**) IL17a, and (**h**) FoxP3 expression was also determined from CD69+CD4+ T cells. N=7. Data points represent individual mice, means ± SD are shown. For (b)-(h), One-way ANOVA with Tukey’s multiple comparisons: *p<0.05; **p<0.01; ***p<0.001; ****p<0.0001; ns = not significant.

### Ad(Spike) cross-reactive antibodies persist and are not affected by BCG pre-immunization

One goal of vaccine development is to promote heterologous reactivity. This is particularly the case with the rapidly evolving SARS-CoV-2 virus where ongoing emergence of novel variants is an important and continuing public health concern. Thus, we assessed if the administration of BCG affected the ability of Spike antibodies generated by an i.n. Ad(Spike) vaccine based on the ancestral (Wuhan) sequence to subsequently recognize epitopes from SARS-CoV-2 alpha (B.1.1.7), beta (B.1.351), gamma (P.1), delta (B.1.617.2), and omicron variants (B.1.1.529, and BA.2). Six months post-vaccination, serum IgG titres against the ancestral S-protein within Ad(Spike)- and BCG(e)+Ad(Spike)-vaccinated animals were not significantly different between the two vaccinated groups (Figure 6a). To ensure BCG pre-immunization does not affect the cross-reactivity of these antibodies, we conducted ELISAs against S-proteins from variant strains of SARS-CoV-2. We found no significant reduction in the antibody binding capacity of serum from vaccinated animals that were vaccinated with Ad(Spike) (Figure 6b) or pre-immunized with BCG and then vaccinated with Ad(Spike) (Figure 6c) against any of the SARS-CoV-2 strains. These results confirm that BCG administration does not affect the generation of cross-reactive anti-Spike antibodies.

**Figure 6.**
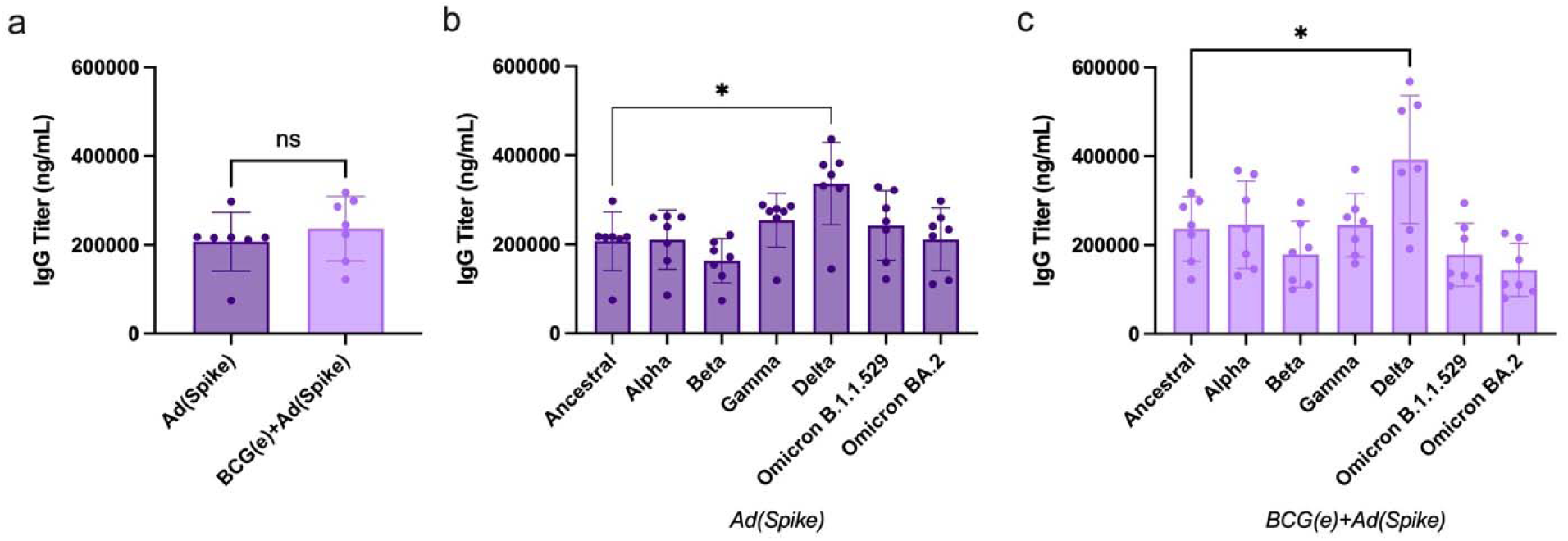
Production of cross-reactive antibodies to variant spike proteins is not altered by BCG. Spike-specific IgG was calculated from serum from vaccinated animals at 6 months post vaccination by ELISA. (**a**) Antibodies are shown against Spike Wuhan (Ancestral) strain. Cross-reactive antibodies from (**b**) Ad(Spike) vaccinated and (**c**) BCG pre-immunized and Ad(Spike) vaccinated animals were then assessed and compared to the ancestral strain. Antibodies binding strains B.1.1.7 (alpha), B.1.351 (beta), P.1 (gamma), B.1.617.2 (delta), B.1.1.529 (omicron), and BA.2 (omicron) were assessed. N=7. Data points represent individual mice, means ± SD are shown. For (a), (b), Kruskal-Wallis test with Dunn’s multiple comparisons: *p<0.05; ns = not significant. For (c), One-way ANOVA with Tukey’s multiple comparisons: *p<0.05; ns = not significant.

## DISCUSSION

A single intranasal dose of a human adenovirus vectored vaccine has been shown to be sufficient to protect mice from SARS-CoV-2 infection and severe disease^32^. However, the durability of protection from these vaccines remains to be established as most studies utilize short-term challenge models. Using an AdV vectored vaccine that expresses the Spike protein of the Wuhan isolate, and then challenging with the Beta variant of SARS-CoV-2, we demonstrated that a single i.n. dose of a human AdV-based vaccine could cross-protect mice from severe disease months after immunization. Specifically, our Ad(Spike) vaccine successfully prevented mice from developing severe pulmonary pathology when challenged with SARS-CoV-2 2- and 6-months post-vaccination. However, the ability of vaccinated mice to limit initial infection was diminished by 6 months post-immunization, with viral shedding in oral swabs greater than earlier challenge experiments, confirming what was observed in a recent meta-analysis^6^. Thus, we aimed to develop a strategy to prolong the mucosal immunity provided by i.n. administration of Ad(Spike) by harnessing the non-specific effects of BCG^33,34^. Here, we demonstrate that BCG improved the long-term protection and viral control conferred by a single i.n. Ad(Spike) by potentiating cellular, rather than humoral, immunity against the Spike antigen in the lungs.

SARS-CoV-2 continues to cause morbidity and mortality, nearly three years after its emergence in December 2019, due to factors such as variants and breakthrough infections caused by waning immunity from vaccines and/or natural infection. In fact, data show that despite short-term efficacy, antibody levels associated with the current COVID-19 vaccines wane over time^35,36^ and is insufficient to completely prevent infection and transmission^5^. Thus, improved vaccines and novel routes that increase mucosal immunity^12,37^ are needed. Indeed, several groups have demonstrated that when adenovirus vectored SARS-CoV-2 vaccines are administered intranasally, a single immunization is sufficient to confer protection from infection in naïve animals^38,39^. However, clinical trials have not yet demonstrated that i.n. AdV vectored vaccines elicit sufficient or lasting immunity to prevent SARS-CoV-2 replication and transmission^13^. As such, safe immune strategies aiming at potentiating these mucosal responses are currently being explored. One promising option, investigated herein, is based on the ability of the BCG vaccine to act as an indirect promoter of cellular immunity. For many years, BCG has been approved for use in infants across the globe to protect against severe, disseminated forms of TB. As such, BCG is commonly reported as the world’s most widely used vaccine and has an excellent safety record. However, BCG has also recently been shown to reduce mortality due to unrelated infectious agents, including several viruses, as a result of its non-specific, immune-enhancing effects^33,40^. Here we sought to investigate the effect of BCG administration on the efficacy and durability of an i.n. human AdV vectored SARS-CoV-2 vaccine. Overall, we found that pre-immunization with BCG can non-specifically rescue waning immunity from a single-dose, i.n. SARS-CoV-2 vaccine by potentiating local vaccine-specific cell mediated responses without hindering humoral immunity.

The possible contribution of widespread BCG vaccination to SARS-CoV-2 protection in human populations is still unclear; yet, there is growing evidence that BCG vaccination provides ‘trained immunity’ to innate mechanisms^40–42^, which can have antagonistic effects on other unrelated pathogens^23,34,43,44^. This is seen in the curious protection from neonatal sepsis and respiratory infections conferred to BCG immunized babies^19^. It has since been demonstrated that due to intrinsic pathogen associated molecular pattern activation of toll like receptors, BCG induces the activation and reprogramming of monocytes^29,40,41^, resulting in increased expression of cell surface markers and the production of pro-inflammatory cytokines and IFN□ in response to antigenic stimulation^45^. These data contributed to the hypothesis that BCG may provide heterologous potentiation of antigen-presenting cell (APC) function to improve the immunogenicity and efficacy of vaccines, as made evident in human studies of neonatal vaccinations^46,47^ and adult influenza vaccination^48^. In fact, BCG is a natural adjuvant that is at least partly a result of its modified peptidoglycan structure and unusual cell-wall lipid composition^49^ and has been exploited in several novel vaccine efforts including against SARS- CoV-2^28,50,51^. It is particularly impressive that BCG has the potential to deliver benefit to vaccination strategies even when not directly co-administered. Nevertheless, BCG alone does not seem to provide direct protection from SARS-CoV-2. While an early study in humans suggested protective efficacy against SARS-CoV-2 from BCG vaccination^52^, a recent case study demonstrated that this protection was not attributable to BCG over the course of the pandemic due to an underreporting of COVID-19 cases in regions with high BCG-vaccination rates^27,53^. Another group found that BCG alone, failed to provide protection from SARS-CoV-2 in mice and hamsters^23^, although this may depend upon the route of administration. Nonetheless, our data support the idea that BCG alone (nor when combined with an empty AdV vaccine vector) does not confer significant protection against SARS-CoV-2 challenge in a mouse model of infection.

As noted above, data show that despite short-term efficacy, protection from SARS-CoV- 2 infection wanes over time in humans^35,36^. Consistent with this, we also observed an attenuation of protection by 6 months post-immunization. Although prior exposure to BCG did not change Ad(Spike) protection 2 months post-vaccination (Supplemental Figure 2), when mice were primed with BCG and then vaccinated with Ad(Spike), significant protection was maintained for at least 6 months post-vaccination. Due to the numerous possible effects of BCG on the innate and adaptive immune responses, we chose to focus on its impact on the type of humoral immunity generated by a single-dose Ad(Spike) vaccine by first assessing the quality and quantity of serum and BALF antibodies. We observed that regardless of their exposure to BCG, mice that were vaccinated with Ad(Spike) displayed robust serum and BALF Spike-specific antibodies, both IgG and IgA which, while slowly declining, were maintained at high titres to at least 6 months post-vaccination. These antibodies maintained high affinity to the receptor binding domain of the Spike protein, confirming that they were capable of neutralizing viral entry. Reassuringly, these antibodies also displayed cross-reactivity against all Spike variants tested, including the beta variant, which was used in our challenge study. However, despite these seemingly high titres of neutralizing IgG1/2c and IgA antibodies, viral RNA and, to a lesser degree, live viral particles remained high in the absence of BCG. Thus, we hypothesized that the bacilli promoted cellular rather than humoral immunity in BCG(e)+Ad(Spike)-immunized mice.

Our antigen-specific assay demonstrated that although Ad(Spike) alone showed a trend towards increased expression of cytotoxic activity in antigen-primed T cells, these differences became significant only in mice that were pre-immunized with BCG. Indeed, BCG has been shown to enhance intramuscular vaccine-induced circulating Spike-specific CD4^+^ T cells in humans^25^, and our results reveal that it can also potentiate the generation of antigen-specific CD4^+^ T cells in tissues. Many reports suggest BCG acts by training monocytes to have higher expression of MHC-II, CD80, and CD86, enhancing antigen presentation to T cells^54,55^. Concomitantly, adaptive immune responses after BCG vaccination also involves the activation of CD8+ T cells^34^. Here, we observed a clear potentiation of Spike-specific cytotoxic CD8+ T cells up to at least 6 months after vaccination. Furthermore, cytokines secreted by BCG-exposed monocytes, such as IL1β and IL6, are key contributors to CD4^+^ T-cell differentiation into Th1 and Th17 subsets^56,57^ which is consistent with the increased CD4^+^ T-cell expression of IFN□ or IL17a we observed in BCG-immunized mice even in the absence of Ad(Spike). Thus, likely, our assay could not distinguish Spike-specific from bystander Th1 and Th17 cells generated prior to BCG administration.

Our study used wild-type C57BL/6 mice, which necessitated using a SARS-CoV-2 strain that binds the mouse ACE-2 (mACE-2) receptor. For this reason, we challenged the C57BL/6 mice with the B.1.351 strain, capable of binding mACE-2 and establishing infection^58^. Thus, we demonstrated cross-protection delivered by our ancestral strain-based AdV vaccine in a context of SARS-CoV-2 infection, which resembles human disease^59^. However, these mice are not typically used for SARS-CoV-2 vaccine studies, as C57BL/6 mice do not display all the hallmark features of severe pathology and typically recover from infection^60^. As follows, the lung pathology observed in our model was modest, posing a limitation in our ability to discriminate severe disease between vaccinated groups. Furthermore, although lung-cell memory responses were increased in BCG pre-immunized animals, our long-term infection model of 6 months post-vaccination may have been insufficient to clearly assess the synergistic effects of BCG on Ad(Spike). We propose that the roles these memory responses play may become more obvious in longer-term studies when protection from Ad(Spike) alone is further reduced.

Despite the limitations of our study, we offer insight into the effects of prior BCG immunization on reinforcing vaccine efficacy in a mouse model of SARS-CoV-2. Interestingly, due to the long-term persistence of viable BCG in our experimental design (Supplemental Figure 4), this study also raises the issue of vaccine efficacy in the context of other persistent bacterial co-infections, as well as components of the normal mucosal microbiota, and represents an important future line of inquiry. As preclinical vaccine testing is conducted in naïve animals, the role of immunological memory from previous vaccination and persisting infections should be addressed in relation to vaccine efficacy.

Collectively, our results present a novel vaccination approach that can potentially curb viral pandemics by potentiating the long-term efficacy of a next-generation adenovirus-vectored mucosal vaccine when provided in the context of the safe and widely distributed BCG vaccine. This approach can also have an impact in other mucosal and non-mucosal vaccination strategies as well. Going forward, these strategies also may provide a viable solution to ensure a more rapid and equitable distribution of vaccines among developing nations where BCG is already firmly entrenched within many vaccination programs. In addition, the combination with BCG also may serve to alleviate concerns over the safety of adenoviral-based vaccines, as it may allow for a reduced dosing schedule or a reduction in the viral titre required to achieve effective vaccination.

## MATERIALS AND METHODS

### Study design

The primary objective of this study was to determine the effect of BCG on subsequent vaccination with a recombinant SARS-CoV-2 S-protein expressing adenoviral vectored vaccine. Experimental units are defined as individual animals. Sample sizes were empirically estimated based on previous data considering the anticipated variation of the results and statistical power needed, while also minimizing the number of animals used. C57BL/6 mice were randomly attributed to treatment groups. To minimise potential confounders, mice were matched for age and sex. Blinding: For all challenge experiments, staff performing infections and sample harvesting were blinded to the different groups and were only unblinded after data analysis. Inclusion/Exclusion: Aside from a small number of deaths in one of the control groups (Ad(e)+BCG(e)), no other animals were excluded from the analysis.

### Animal ethics

All animal procedures were performed in accordance with the Institutional Animal Care and Use Guidelines approved by the Animal Care and Use Committee at McGill University (Animal Use Protocol 8190). Mouse housing, husbandry, and environmental enrichment can be found within the McGill standard operating procedures (SOP) #502, #508, and #509. Animals were monitored for adverse events for 3 days post-vaccination and weekly until the end of each experiment. Humane intervention points were monitored according to McGill SOP #410. Challenge experiments were performed in compliance with the Canadian Council on Animal Care guidelines and approved by the Animal Care Ethics Committee at the University of Toronto (APR-00005433-v0002-0). All animals were humanely sacrificed at endpoint by anaesthesia with isoflurane before euthanasia by carbon dioxide asphyxiation, followed by pneumothorax and blood collection by cardiac puncture.

### Cell lines and reagents

Cell lines were obtained from commercial sources, passed quality control procedures, and were certified and validated by the manufacturer. SF-BMAd-R cells were validated for identity, as human derived^61^. All reagents were validated by the manufacturer or has been cited previously in the literature. When available, RRID tags have been listed in the text and in the reagent repository (Supplemental Table 2).

### *Mycobacterium bovis* BCG Danish culture

*Mycobacterium tuberculosis* variant *bovis* BCG (ATCC 35733), provided to us by Dr. Marcel Behr (Research Institute of the McGill University Health Centre), was grown in Middlebrook 7H9 broth (BD, Mississauga, ON, Canada) supplemented with 10% ADC (8.1g/l NaCl, 50g/l BSA Fraction V (Millipore Sigma, Billerica, MA, USA), 20g/l glucose), 0.2% glycerol and 0.05% Tween 80, or on Middlebrook 7H11 agar (BD) supplemented with 10% OADC enrichment (as per ADC plus 0.6ml/l oleic acid, 3.6mM NaOH). The BCG-Danish strain used in these experiments was transformed with an empty pMV361(hygromycin^R^) plasmid and was initially selected in the presence of 50μg/ml hygromycin (Wisent, Saint-Jean-Baptiste, QC, Canada). This plasmid was integrated into the non-essential L5 phage attachment site (*attB*) located within the BCG chromosome. We termed this strain BCG(e).

### Preparation of *M. bovis* BCG-Danish cultures for mouse immunization

100 ml BCG cultures were grown in 7H9/ADC medium to an OD_600nm_ = 0.6. After spinning at 3000 rpm, cells were washed twice with PBS containing 0.05% Tween-80 (PBS-Tw) and resuspended in 6 ml of this buffer. After passing 10 times through a 22G x1” and 10 times through a 27G x1/2” needle, the suspension was mixed with 4 ml of sterile 50% glycerol in PBS. Aliquots were made and frozen at -80^0^C. Prior to immunization, an aliquot was thawed, and 10- fold serial dilutions were plated on 7H11/OADC+ HYG for quantification, yielding a value of ∼1.5 x10^8^ cfu/ml. On the day of the immunization, these glycerol stocks were diluted (1/20) in PBS-Tw and 200 ul (containing ∼ 1.5 x10^6^ cfu) were injected i.p. into the lower right quadrant of the abdomen of the mice using a 28Gx1/2” needle and an insulin syringe. Inocula were quantified by plating 10-fold serial dilutions on 7H11/OADC+ HYG.

### Generation of Ad(Spike) vector

The AdSpike construct was developed following a similar protocol as described^62^. Briefly, the Spike gene cassette combined a Kozak sequence with the full length of the Spike protein (Genbank accession number QHU36824.1), codon optimized to mouse and human expression avoiding restriction sites Bgl2, Pac1, and Pme1, followed by a Kpn1 restriction site and the poly-A signal “TCTAGACTCGACCTCTGGCTAATAAAGGAAATTTATTTTCATTGCAATAGTGTGTTGGAATTTTTTGTGTCTCTCACTCGGAAGGACATATGGGAGGGCAAATCATTTGCGGCCGCGATATC” (GenScript, Piscataway, NJ, USA). The gene cassette was flanked by Bgl2 sites and synthesized by Integrated DNA Technologies (Coralville, IA, USA) then cloned into the vector, pShuttle-CMV-Cuo^63^. Primers to confirm gene sequence can be found in Supplemental Table 3. The plasmid containing our recombinant non-replicating human adenovirus serotype 5 (E1 and E3 genes removed (ΔE1-, ΔE3-); 1^st^ generation) encoding the S-protein gene was made through homologous recombination in AdEasier-1 cells (strain), a gift from Dr. Bert Vogelstein (Addgene plasmid #16399) (Addgene, Watertown, MA, USA) ^64^. It was then linearized with *PacI* and transfected into HEK293A cells (RRID:CVCL_6910). Our recombinant adenovirus was then amplified using SF-BMAd-R cells ^61^ in 3 batches (Ad(Spike) 1-3), combined, and purified by ultracentrifugation on CsCl gradients as described previously^65^, before titration using a TCID_50_ assay. A second human adenovirus serotype 5 (ΔE1-, ΔE3-; 1^st^ generation), lacking a gene cassette, was used as a negative control.

### Western blot assays

To determine protein expression by Ad(Spike), cell lysates of Ad(Spike) infected HEK293A cells were assessed. Briefly, cells were infected at a multiplicity of infection of 5 particles per cell and incubated for 48-72 hours, pelleted, and then lysed (0.1M Tris, 10 μL EGTA, 50 μL Triton-100, 0.1M NaCl, 1mM EDTA, 25 μL 10% NaDeoxycholate, 1X protease inhibitor, in ddH_2_O). Cell lysates were then resolved on an SDS-PAGE gel under reducing conditions followed by transfer onto a nitrocellulose membrane. The membrane was subsequently blocked in phosphate buffered saline (PBS) with 0.05% Tween 20 (PBS-T) and 5% milk (Smucker Foods of Canada Corp, Markham, ON, Canada) (PBS-TM). The membrane was then incubated with rabbit anti-SARS-CoV-2 Covid-19 Spike RBD coronavirus polyclonal antibody (RRID:AB_258251) diluted 1:5,000 in PBS-TM overnight at 4°C. The membrane was then washed in PBS-T before incubation with horseradish peroxidase (HRP)-conjugated anti-rabbit IgG (IgG-HRP) (Rockland Immunochemicals, Pottstown, PA, USA) diluted 1:10,000 in PBS-T for one hour at room temperature. After incubation the membrane was washed again and developed using SuperSignal West Pico Plus Chemiluminescent Substrate (ThermoFisher Scientific, Waltham, MA, USA).

### Protein expression and purification

SARS-CoV-2 Spike variants Wuhan, B.1.1.7 (alpha), B.1.351 (beta), P.1 (gamma), B.1.617.2 (delta), B.1.1.529 (omicron), and the RBD portion of the Wuhan Spike variant were obtained from the National Research Council of Canada. Recombinant, “tagless” Spike proteins were produced as previously described^66^. Recombinant RBD protein was produced as previously described^67,68^. Spike variant BA.2 (omicron) was obtained through BEI Resources, NIAID, NIH: Spike Glycoprotein (Stabilized) from SARS-Related Coronavirus 2, BA.2 Lineage (Omicron Variant) with C-Terminal Histidine and Avi Tags, Recombinant from HEK293 Cells, NR-56517.

### Immunization and challenge protocol in mice

Six- to eight-week-old female C57BL/6 mice were ordered from Charles River Laboratories (RRID:IMSR_CRL:027) (Senneville, QC, Canada). Each mouse was immunized at weeks 0 and 4 by intraperitoneal (i.p.) injection of 200μL of BCG(e) and i.n. administration of 30μL of adenovirus formulations, respectively. Group 2 was removed from the long-term challenge experiment since negative control animals displayed similar viremia to the PBS control in the short-term challenge model. See Table 1 for more precise group descriptions. Mice were bled from the saphenous vein at weeks 0, 4, 8, 12, and 18. Mice immunized for humoral and cell-mediated immunity assessment (n=6) were euthanized 6 months after the final immunization and blood, spleens, and lungs were collected. Mice immunized for challenge studies (n=12) were transferred to the University of Toronto and challenged with 10^6^ TCID_50_ of SARS-CoV-2 South African strain (B.1.351) 2 (n=6) or 6 months (n=6) post-Ad(Spike)-vaccination. TCID_50_ was determined using the Spearman–Kärber method^69^. Oral swabs were taken from mice on days 1, 3, and 5 post-challenge in DMEM. Mice were euthanized 5 days after challenge and lungs were collected.

**Table 1.** Animal groups

| Group | Prime (i.p.) | Boost (i.n.) |
| --- | --- | --- |
| 1: PBS | PBS | PBS |
| 2: BCG(e)+Ad(e) | 10<sup>6</sup> colony forming units (cfu) of BCG (Danish strain) carrying an integrated, empty pMV361(Hygromycin<sup>R</sup>) plasmid; referred to here as BCG(e) | 10<sup>9</sup> TCID<sub>50</sub> of an empty, non-replicating human adenovirus serotype 5 vector; referred to here as Ad(e) |
| 3: Ad(Spike) | - | 10<sup>9</sup> TCID<sub>50</sub> of recombinant AdV expressing the full-length spike (S)-protein; referred to here as Ad(Spike) |
| 4: BCG(e)+Ad(Spike) | 10<sup>6</sup> cfu of BCG(e) | 10<sup>9</sup> TCID<sub>50</sub> of Ad(Spike) |

### Quantification of viral load

Quantities of infectious virus was determined by determining the median tissue-culture infectivity dose (TCID_50_) using methods that have been described previously^70^. Briefly, Vero E6 cells were seeded into plates and incubated overnight at 37°C. On the following day, media was removed, and samples were added and serially diluted using ten-fold dilutions. Plates were incubated at 37°C for 1□h. After incubation, the media was removed and replaced with complete DMEM, and plates were incubated at 37°C for 5 days. Cells were examined for cytopathic effect (CPE) at 5 dpi. TCID_50_ was defined using the Spearman–Kärber method^69^.

### qRT-PCR

Viral RNA loads were calculated as previously described^70^. Briefly, viral RNA was extracted using the QIAamp viral RNA kit (Qiagen, Hilden, Germany) according to the manufacturer’s guidelines. SARS-CoV-2 viral RNA detection and quantification was performed using the Luna Universal Probe One-Step RT-qPCR kit (New England Biolabs, Whitby, ON, Canada) on the Rotor-gene Q platform (Qiagen). For quantification, standard curves were generated using a synthetic plasmid containing a segment of the E-gene (GenScript) and interpolation was performed as described by Feld et al.^71^. The limit of quantification was determined to be 20 copies/mL.

### Spike and RBD-specific IgG, IgG1, IgG2c, IgA quantification and IgG avidity assays

Briefly, high binding 96-well plates (Greiner Bio-One, Frickenhausen, Germany) were coated with recombinant Spike or RBD (0.5 μg/mL) in 100 mM bicarbonate/carbonate buffer (pH 9.6) along with various standard curves (IgG, IgG1, IgG2c, IgA: serially diluted from 2000 ng/mL to 1.953 ng/mL) overnight at 4°C. Then, plates were blocked with 2% bovine serum albumin (BSA; Sigma Aldrich, St. Louis, MO, USA) in PBS-T (blocking buffer) for 1 hour at 37°C before samples diluted in blocking buffer were added in duplicate. Nasal wash samples were run in singlet. When running serum for total Spike/RBD-IgG, an additional set of serum samples was run to determine IgG avidity. Plates were incubated for 1 hour at 37°C then washed with PBS (pH 7.4). For IgG avidity assessment, the additional set of samples received 8M urea, while blocking buffer was added to the first set and the standard curve. Plates were covered and incubated for 15 minutes at room temperature protected from light, washed 4 times, and then blocked again with blocking buffer for 1 hour at 37°C. Next, plates were washed with PBS and anti-mouse IgG-HRP (Sigma Aldrich) was diluted 1:20,000 in blocking buffer and applied for 30 minutes at 37°C. For other immunoglobulins, the same protocol was followed without the additional avidity steps and the appropriate HRP-conjugated antibody was applied. Both IgG1- and IgG2c-HRP were diluted 1:20,000 in blocking buffer and applied for 30 minutes at 37°C. For IgA, HRP-conjugated anti-mouse IgA (Sigma Aldrich) was diluted 1:2,000 in blocking buffer and applied for 1 hour at 37°C. Plates were washed a final time with PBS and 3,3’,5,5’- Tetramethyl benzidine (TMB) substrate (Sigma Aldrich) was added to each well. The reaction was stopped after 15 minutes using H_2_SO_4_ (0.5M; Fisher Scientific, Waltham, MA, USA) and the optical density (OD) was measured at 450 nm with an EL800 microplate reader (BioTek Instruments Inc., Winooski, VT, USA). Concentrations of Spike/RBD specific antibodies were calculated by extrapolation from respective standard curves and multiplied by the dilution factor. IgG avidity indices were calculated by dividing the IgG titre in the urea conditions by the IgG titre in the non-treated condition.

### Surrogate virus neutralization test

Neutralizing antibodies were assessed using the cPass SARS-CoV-2 Neutralization Antibody Detection Kit (GenScript) according to manufacturer’s instructions with the following changes: To collect semi-quantitative results, the kit was run with the SARS-CoV-2 Neutralizing Antibody Calibrator to create a standard curve used to determine the concentration of neutralizing antibodies. BALF samples were run neat or diluted (1:3) and serum samples were diluted (1:150). Samples that gave values above the 30% signal inhibition cut-off value were multiplied by the dilution factor and reported as Units/mL. Data reported according to the manufacturer’s guidelines can be found in Supplemental Tables 4 (serum) and 5 (BALF).

### BALF and lung collection

Six months after the last immunization, unchallenged mice were euthanized and the lungs were collected. Bronchoalveolar lavage fluid (BALF) was collected by combining four lung washes of 0.5mL of cold PBS+protease inhibitor. Lungs were collected in 1mL cold RPMI. Lungs were digested enzymatically for 30 minutes at 37°C and 5% CO_2_ with a cocktail of DNase I (200 µg/mL, Sigma Aldrich), LiberaseTM (100 µg/mL, Roche, Indianapolis, IN, USA), hyaluronidase 1a (1 mg/mL, Life Technologies, Carlsbad, CA, USA), and collagenase XI (250µg/ml; Life Technologies) in RPMI-1640 as described previously^72^. Cells were then washed with RPMI- 1640 media containing 1% Penicillin/Streptomycin and 5% FBS. Sterile, filtered ammonium-chloride-potassium (ACK) buffer was used to lyse red blood cells. Filtration through a 0.7 μM strainer was performed and the remaining viable cells were recovered.

### Quantification of cytokine-secretion in T cells by multi-parametric flow cytometry

Lung lymphocytes were seeded into 96-well flat bottom plates (BD) at 10^6^ cells in 200 uL/well. Duplicate cultures were stimulated with or without a combined preparation of Peptivator Peptide Pools of the complete Spike protein and predicted immunodominant sequences (Miltenyi Biotec, Bergisch Gladbach, North Rhine-Westphalia, Germany) in RPMI (0.3 μg/mL final concentration) for 18 and 96 hours at 37°C + 5% CO_2_. For the last 6 hours of incubation, protein transport inhibitor was prepared according to the manufacturer’s guidelines (RRID:AB_2869014, BD Science, San Jose, CA, USA) and added to all samples. Cells stimulated with phorbol 12-myristate 13-acetate (Thermofisher Scientific) and ionomycin (Thermofisher Scientific) were processed as positive controls. All staining and fixation steps took place at 4°C protected from light. Briefly, the cells were washed twice with cold PBS and stained with 50μL/well fixable viability dye eFluor 780 (Thermofisher Scientific) diluted at 1:1000 for 20 minutes. Cells were washed once with PBS. All surface stains were diluted 1:50 in PBS and 50μL/well of extracellular cocktail was applied for 30 minutes. The following antibodies made up the extracellular cocktail: CD3-BUV395 (145-2C11, RRID: AB_27382, BD Biosciences, Franklin Lakes, NJ, USA), CD4-AF700 (RM4-5, RRID: AB_49370, BioLegend, San Diego, CA, USA), CD8b-BV510 (H35-17.2, RRID: AB_2739908, BD Biosciences) and CD69-FITC (H1.2F3, RRID: AB_313108, BioLegend). Cells were then washed as before and fixed with the eBioscience FoxP3 transcription factor staining buffer (Thermofisher Scientific) overnight. The next day, plates were washed with 1X permeabilization buffer (perm buffer) (Thermofisher Scientific) and stained with an intracellular cocktail of antibodies diluted 1:50 in perm buffer applied as 50μL/well for 30 minutes. The intracellular cocktail was made up of: FOXP3-Pe-Cy7 (FJK-16s, RRID: AB_891552, Thermofisher Scientific), IFNγ-BUV737 (XMG1.2, RRID: AB_2870098, BD Biosciences), IL-17A-APC (TC11-18H10.1, RRID: AB_536018, BioLegend), IL-4-BV421 (11B11, RRID: AB_2562594, BioLegend), GrB-PE (QA1602, RRID: AB_2687032, BioLegend), and TNFα-PerCP-Cy5.5 (MP6-XT22, RRID: AB_961434, BioLegend). After staining, cells were washed twice with perm buffer and resuspended in PBS 1X and acquired on a BD LSRFortessa X-20 (BD Science). Flow data were analysed using FlowJo software (version 10.0.8r1) (Treestar, Ashland, OR, USA). Our gating strategy is shown in Supplemental Figure 5.

### Histological analysis

Organs collected from the challenged animals at the time of necropsy were placed in 10% phosphate-buffered formalin. Collected tissues were subsequently processed for histopathology, and slides were stained with haematoxylin and eosin (H & E), to assess tissue architecture and inflammation, and Masson’s trichrome, to determine the progression of fibrosis. Sections of lungs were examined and scored by a pathologist who was blinded to the experimental groups. Lungs were evaluated for fibrosis, the presence or absence of features of cell or tissue damage (CTD: necrosis of bronchiolar epithelial cells (BEC), inflammatory cells and/or cellular debris in bronchi, intraepithelial neutrophils, alveolar emphysema), circulatory changes and vascular lesions (CVL: alveolar hemorrhage, significant alveolar edema, vasculitis/vascular endothelialitis), reactive inflammatory patterns (RIP: necrosuppurative bronchitis, intraalveolar neutrophils, and macrophages, mononuclear infiltrates around airways, presence of polymorphonuclear granulocytes, perivascular mononuclear cuffs, and mesothelial reactivity), as well as regeneration and repair (RR: alveolar epithelial hyperplasia/regeneration, BEC hyperplasia/regeneration) as described elsewhere ^70,73^. After the scoring was completed, lung pathology scores were tabulated. Processed and stained lung slides were digitized using Aperio AT2 (Leica Biosystems, Wetzlar, Germany).

### Statistical analysis

Statistical analysis was performed using GraphPad Prism 9 software (La Jolla, CA, USA). Data were assessed for normality using Shapiro-Wilk tests. Non-parametric data were analysed by Kruskal-Wallis tests with Dunn’s multiple comparisons. When appropriate, one-way and two-way ANOVAs were employed with Tukey’s multiple comparisons. P values <0.05 were considered significant.

### List of supplementary materials

Figures S1 to S6

Tables S1 to S5

ARRIVE Checklist

MDAR Reproducibility Checklist

## Supporting information

ARRIVE Checklist

MDAR Checklist

## Acknowledgements

We would like to thank Annie Beauchamp, Rami Karkout, Sarah Santoso, Louis Cyr, Angela Brewer, and Raidan Alyazidi for their contributions during animal sacrifice and advice for experimental procedures as well as the other members of the Ward/Ndao laboratory for their support. We would also like to thank Dr. Marcel Behr for kindly providing us with the Danish strain of BCG. We would like to thank Heather Tyra and Francois Francoeur at IDT for their assistance in making our gene construct. We would like to thank Nazila Nazemi-Moghaddam and Claire Guilbault of the National Research Council of Canada for their support manufacturing, amplifying, and titrating our recombinant adenovirus. We thank the staff from the high-containment lab (Sunnybrook Hospital) for their technical assistance. We would like to thank the members of the Mammalian Cell Expression Section of the NRC-HHT for their contribution to producing and purifying the recombinant proteins and those members at BEI resources for their contributions of recombinant proteins. In addition, we would like to thank the Immunophenotyping Platform at the Research Institute of the McGill University Health Centre (RI-MUHC). We would like to thank Kim Babin and Bruce Lu, from Euroimmun and GenScript, respectively, with their help obtaining cPass neutralizing antibody kits. We would also like to thank Ara Xiii for their help in editing our manuscript. Finally, we would like to thank all the entities which contributed to this work financially including the McGill Interdisciplinary Initiative in Infection and Immunity (MI4) program, the Foundation of the MUHC, the R. Howard Webster Foundation, the Foundation of the Montreal General Hospital, and the RI- MUHC.

## Author contributions

Study design: DJP, PD, MBR, and M Ndao in collaboration with RK, GPK, and RG

Funding acquisition: M Ndao, MBR, MO

Vaccine design: DJP and PD with assistance from SME.

Animal vaccinations and immunogenicity experiments: DJP, CKP, PD

Animal sacrifice, sample collection, and sample processing: DJP, PD, FA, and LL

Histology imaging and scoring: AL and POF

Recombinant protein production: MS and YD Manuscript preparation: DJP, FA

Manuscript editing and contribution: PD, GGB, M Naghibosodat, CAP, RK, MBR, M Ndao

Animal challenge experiments: M Naghibosadat, GGB, and RK

All authors have read and approved the final version of this manuscript.

## Data and materials availability

All data associated with this study are present in the paper or the Supplementary Materials. Requests may be made by contacting the corresponding author.

