## Supplementary Data for "BCG administration promotes the long-term protection afforded by a single-dose intranasal adenovirus-based SARS-CoV-2 vaccine"

**Supplemental Figures**

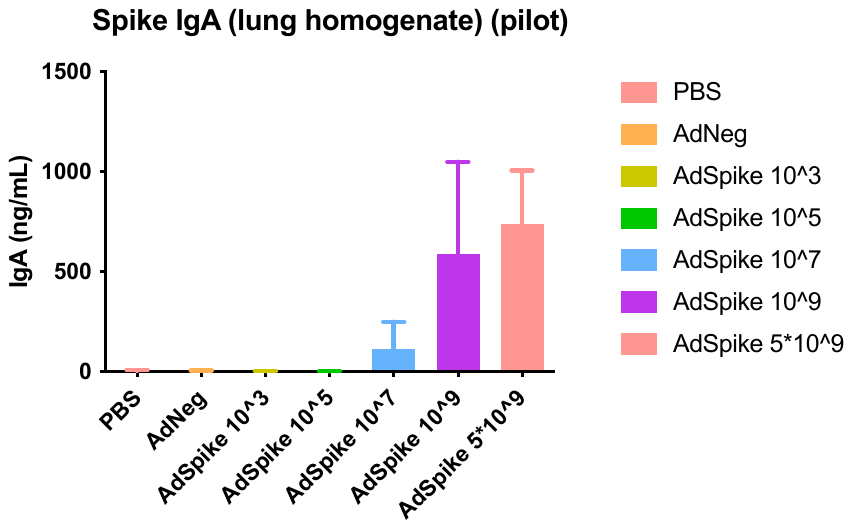

**Supplemental Figure 1. Induction of spike-specific lung IgA based on Ad(Spike) dosage**

A pilot study was run to determine optimal dosage of Ad(Spike) for intranasal administration. Nine weeks after immunization, animals were sacrificed, and spike-specific IgA was calculated in lung homogenates by ELISA. AdNeg=human adenovirus serotype 5 containing an empty gene cassette administered at 10^9^ TCID_50_. All other doses are given in TCID_50_. N=5. Data points represent individual mice, means ± SD are shown.

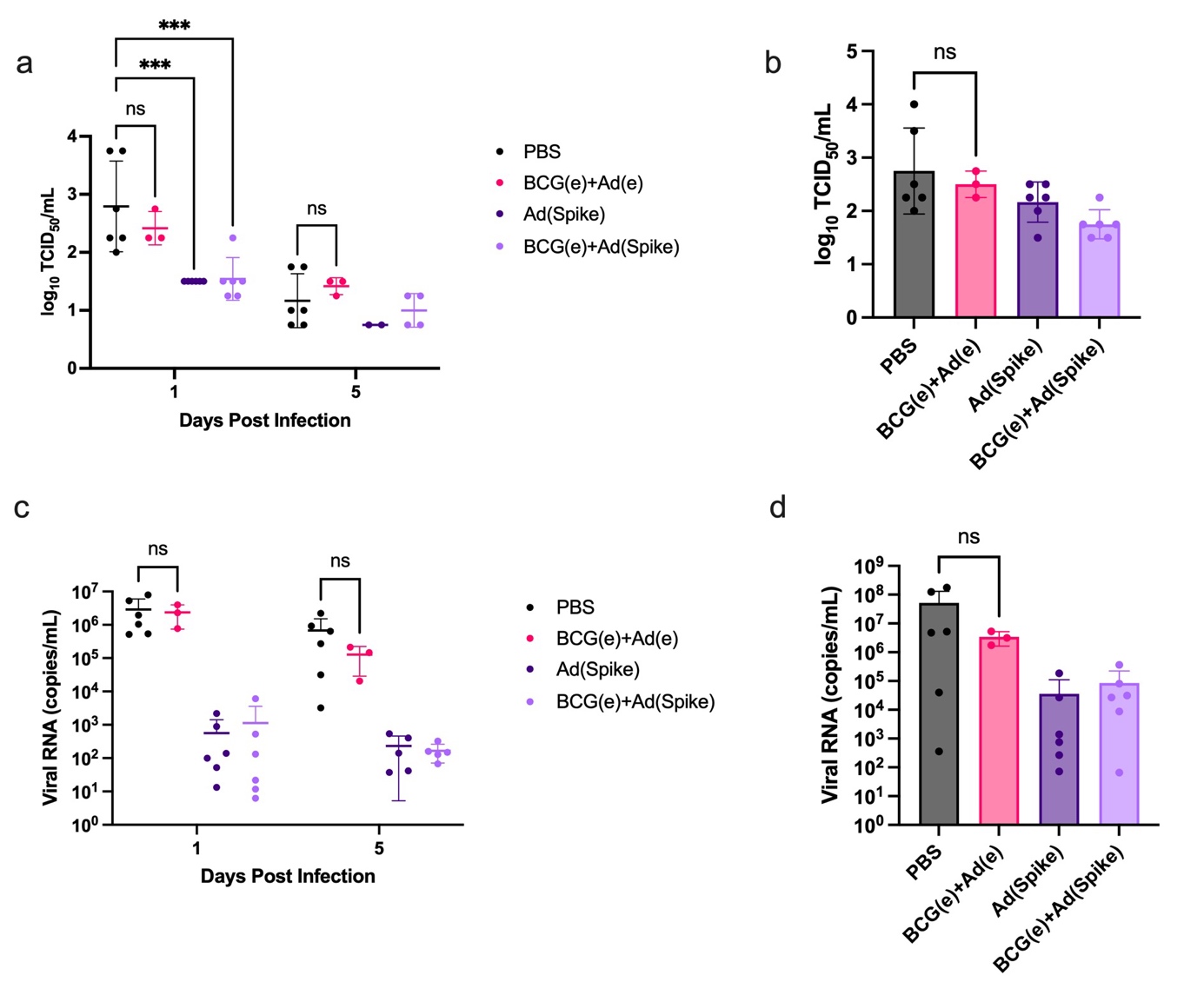

**Supplemental Figure 2. BCG(e) alone does not provide protection from SARS-CoV-2 infection in C57BL/6 mice.**

Mice were pre-immunized with 10^6^ BCG(e) or PBS i.p. at -1 month and immunized with either Ad(e) or Ad(Spike) i.n. at 0 months. Two months post-vaccination animals were infected with 10^6^ TCID_50_ SARS-CoV-2 South African strain (B.1.351). Infectious viral load (TCID_50_) in (a) oral swabs at 1 and 5 dpi, and (b) lungs at 5 dpi, quantified by the Spearman–Kärber method. Viral RNA in (c) oral swabs at 5 dpi and (d) lungs at 5 dpi. N=3-6; note that 3 animals died within the BCG(e)+Ad(e) group post-infection. Data points represent individual mice, means ± SD are shown. For (a), (c), Two-way ANOVA with Tukey’s multiple comparisons: ***p<0.001; ns = not significant. For (b), (d), Kruskal-Wallis test with Dunn’s multiple comparisons: ns = not significant.

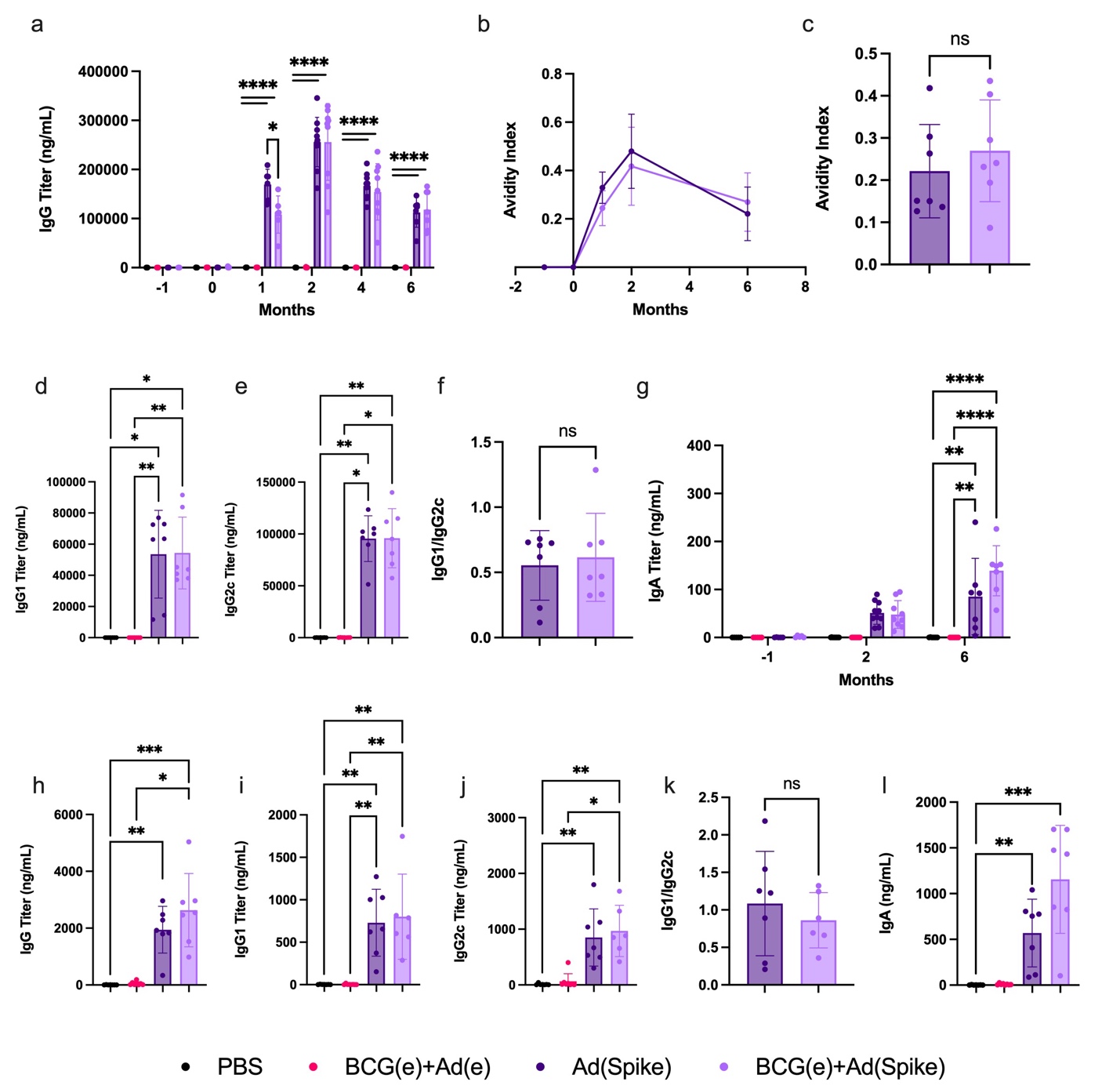

**Supplemental Figure 3. RBD-specific antibody response**

RBD-specific antibodies in the (a)-(g) serum and (h)-(l) bronchoalveolar lavage fluid (BALF). (a) IgG titers in mouse sera throughout the study schedule determined by ELISA. (b) IgG avidity index at -1, 0-, 1-, 2-, and 6-months post vaccination with 6 months shown in (c). (d) IgG1 and (e) IgG2c at 6 months post vaccination. The ratio of RBD-specific IgG1/IgG2c at 6 months post vaccination is given in (f). (g) IgA titers in mouse sera calculated at 0-, 3-, and 6-months post vaccination. N=7-10. Spike-specific (h) IgG, (i) IgG1, (j) IgG2c, (l) IgA in BALF at 6 months post vaccination calculated by ELISA with the ratio of IgG1/IgG2c given in (k). N=7. Data points represent individual mice, means ± SD are shown. For (a), (g), Two-way ANOVA with Tukey’s multiple comparisons: *p<0.05;**p<0.01; ****p<0.0001. For (c), (k), One-way ANOVA with Tukey’s multiple comparisons: ns = not significant. For (d)-(f), (h)-(j), (l), Kruskal-Wallis test with Dunn’s multiple comparisons: *p<0.05; **p<0.01; ***p<0.001; ns = not significant.

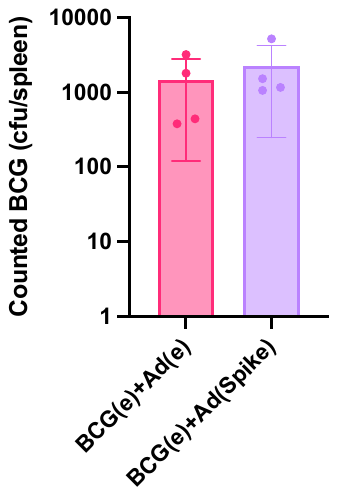

**Supplemental Figure 4. BCG persists throughout long term challenge**

Animals which were pre-immunized with BCG(e) (-1 months) and vaccinated with either Ad(e) or Ad(Spike) (0 months) were euthanized 6 months post-vaccination. Upon necropsy, spleens were isolated, homogenized and then plated on Middlebrook 7H11/OADC for BCG quantification. Both groups show viable BCG present in the spleens. Data is represented as means ± SD with each point being an individual mouse. N=4.

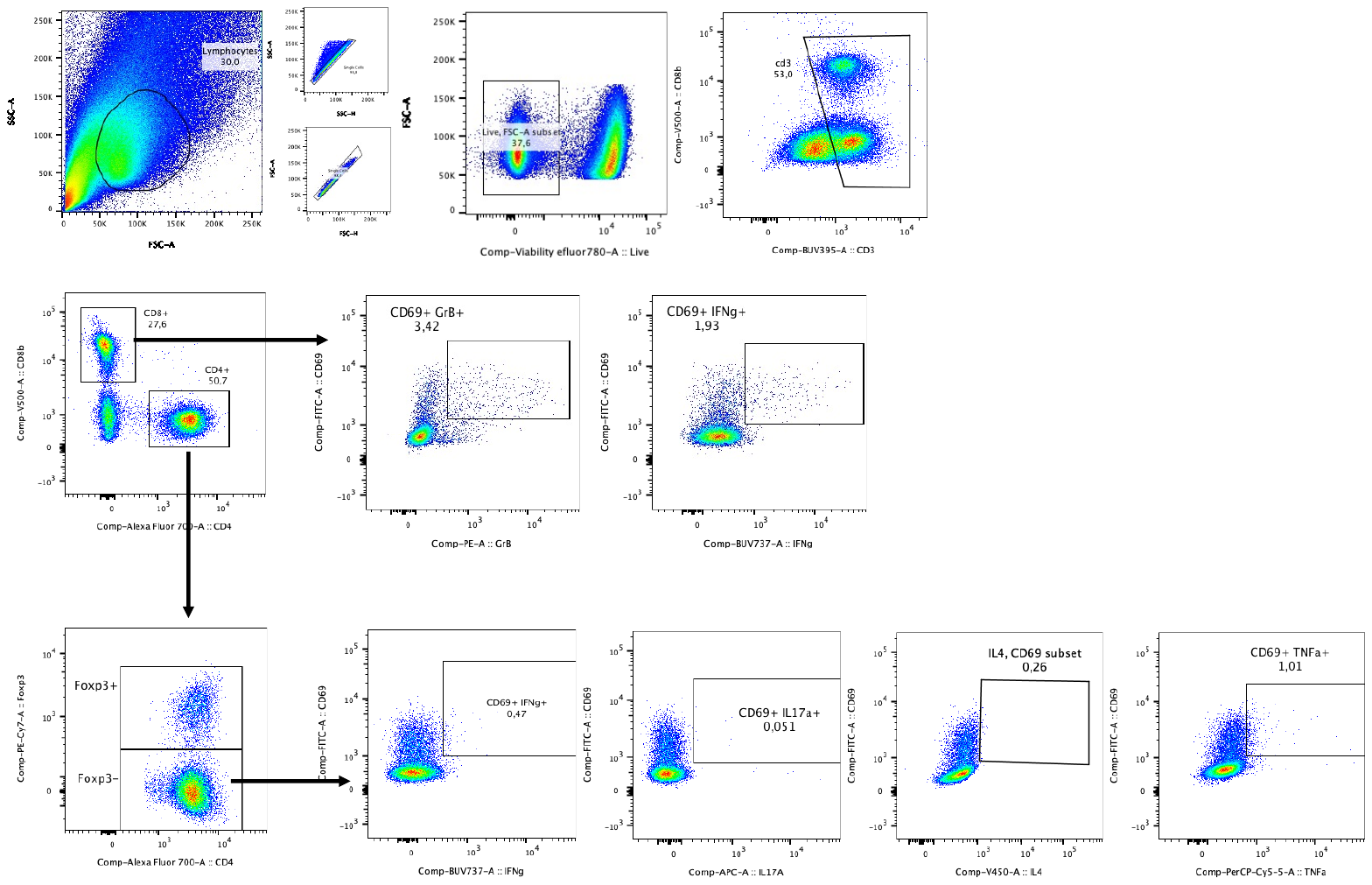

**Supplemental Figure 5. Flow cytometry gating strategy**

Splenocytes were isolated 6 months after immunization with Ad(Spike) before restimulation with a pool of SARS-CoV-2 Spike CD8+ epitopes for 96 hours. During acquisition on a flow cytometer, cells were gated on lymphocytes, and then single cells. The live population was chosen and gated on CD3+ cells. CD4+ and CD8+ T cells were gated and CD8+ T cells were assessed on their expression of CD69+GrB+ and CD69+IFNg+. The CD4+ cells were gated as FoxP3- and assessed on their expression of CD69+IFNg+, CD69+IL17a+, CD69+IL4+, CD69+TNFa+. (IFNg=IFNɣ, TNFa=TNFɑ, GrB=Granzyme B).

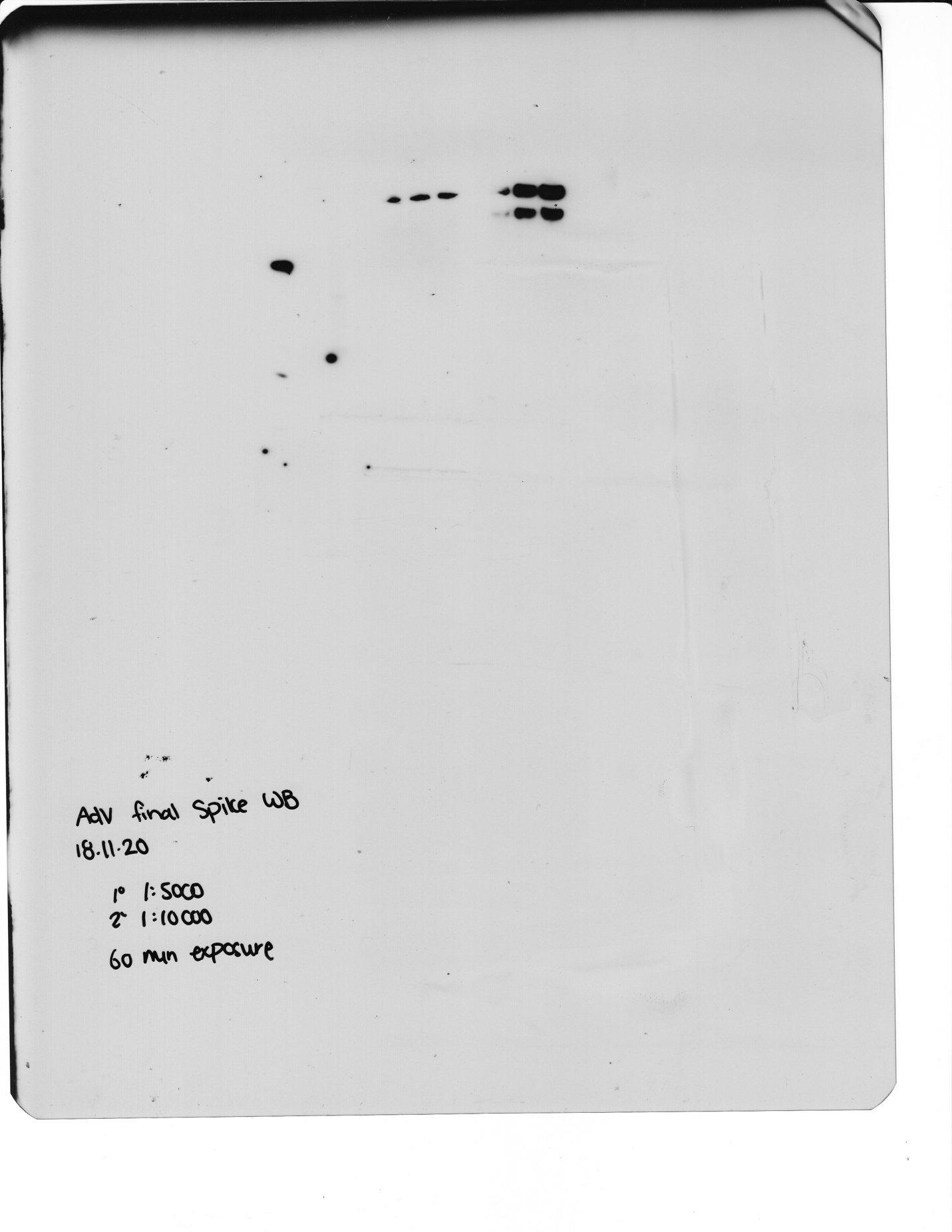

**Supplemental Figure 6. Full Ad(Spike) Western Blot**

Western blot conducted against cell lysates from Ad(Spike) and a second adenovirus, containing a mutated version of the S-protein (not used in this study). This western blot was run with a protein ladder and negative control: adenovirus containing an empty gene cassette, termed Ad(e).

**Supplemental Tables**

**Supplemental Table 1. Observed pathology scoring from immunized and challenged mice in long-term challenge.** Statistics calculated against the PBS control. (Two-way ANOVA with Tukey’s multiple comparisons: *p<0.05)

|  | PBS | Ad(Spike) | BCG(e)+Ad(Spike) |
| --- | --- | --- | --- |
| Fibrosis | 2 | 0 | 0.33 |
| CTD | 6 | 0.33* | 0.17* |
| CVL | 3.33 | 0.17 | 0 |
| RIP | 6.83 | 1.33* | 1.33* |
| RR | 2.83 | 0.17 | 0 |
| Total | 20.99 | 2 | 1.83 |

**Supplemental Table 2. Reagent repository.**

| **Name** | **ID/Catalogue Number (Cat)/RRID** | **Company** | **Validation** |
| --- | --- | --- | --- |
| AdEasier-1 cells | GenBank ID:AY370909, Addgene plasmid #16399 | Addgene, Watertown, MA, USA | A simplified system for generating recombinant adenoviruses. He TC, Zhou S, da Costa LT, Yu J, Kinzler KW, Vogelstein B. Proc Natl Acad Sci U S A. 1998 Mar 3. 95(5):2509-14. 10.1073/pnas.95.5.2509 PubMed 9482916 |
| One Shot Top10 Chemically Competent *E. coli* | Cat:C404010 | ThermoFisher Scientific, Waltham, MA, USA | https://www.thermofisher.com/document-connect/document-connect.html?url=https%3A%2F%2Fassets.thermofisher.com%2FTFS-Assets%2FLSG%2Fmanuals%2Foneshottop10_chemcomp_man.pdf |
| HEK293A cells | RRID:CVCL_6910  Cat: CRL-1573 | American Type Culture Collection (ATCC), Manassas, VA, USA | Graham FL, Smiley J, Russell WC, Nairn R. Characteristics of a human cell line transformed by DNA from human adenovirus type 5. J. Gen. Virol. 1977; 36; 59-74. doi: 10.1099/0022-1317-36-1-59.  https://www.atcc.org/api/pdf/product-sheet?id=CRL-1573 |
| SF-BMAd-R cells | Developed in-lab | National Research Council of Canada, Montreal, QC, Canada | Gilbert R, Guilbault C, Gagnon D, Bernier A, Bourget L, Elahi SM, Kamen A, Massie B. Establishment and validation of new complementing cells for production of E1-deleted adenovirus vectors in serum-free suspension culture. J Virol Methods. 2014 Nov;208:177-88. doi: 10.1016/j.jviromet.2014.08.013. Epub 2014 Aug 23. PMID: 25159033. |
| non-replicating Adenovirus (ΔE1, ΔE3; 1st generation) containing no gene cassette | Produced in-lab | National Research Council of Canada, Montreal, QC, Canada | Garnier, A., J. Côté, I. Nadeau, A. Kamen, and B. Massie. 1994. Scale-up of the adenovirus expression system for the production of recombinant protein in human 293S cells. Cytotechnology 15:145–155. |
| Rabbit anti-SARS-COV2 SARS-COV/SARS-COV-2 Spike RBD Polyclonal Antibody(2019-nCoV) Polyclonal Antibody | Cat: MBS2563840 | MyBioSource, San Diego, CA, USA | https://cdn.mybiosource.com/tds/protocol_manuals/800000-9999999/MBS2563840_TDS.pdf |
| Anti-mouse IgG HRP | Cat:A0168 | Sigma Aldrich | https://www.sigmaaldrich.com/specification-sheets/291/310/A0168-BULK________SIGMA____.pdf |
| SuperSignal West Pico Plus Chemiluminescent Substrate | Cat:34580 | Thermofisher | https://www.thermofisher.com/document-connect/document-connect.html?url=https%3A%2F%2Fassets.thermofisher.com%2FTFS-Assets%2FLSG%2Fmanuals%2FMAN0015920_2162617_SuperSigWestPicoPLUS_Chemil_Substr_UG.pdf |
| Recombinant Spike and RBD Proteins | Produced in-lab |  |  |
| C57BL/6NCrl mice | RRID:IMSR_CRL:027 | Charles River Laboratories, Senneville, QC, Canada | https://www.criver.com/sites/default/files/resources/C57BL6MouseModelInformationSheet.pdf |
| HRP-conjugated anti-mouse IgM | RRID:AB_2794240  Cat:1021-05 | SouthernBiotech, Birmingham, AL, USA | https://resources.southernbiotech.com/techbul/1021.pdf |
| HRP-conjugated anti-mouse IgE | RRID: AB_2868311  Cat:SA5-10263 | Thermofisher | https://www.thermofisher.com/order/genome-database/dataSheetPdf?producttype=antibody&productsubtype=antibody_secondary&productId=SA5-10263&version=216 |
| HRP-conjugated anti-mouse IgA | RRID:  Cat:A4789 | Sigma Aldrich | https://www.sigmaaldrich.com/specification-sheets/510/236/A4789-BULK.pdf |
| TMB Substrate | Cat:T0440 | Sigma Aldrich | https://www.sigmaaldrich.com/specification-sheets/157/638/T0440-BULK________SIGMA____.pdf |
| Goat anti-mouse IgG1-HRP | RRID:AB_2794426  Cat:1071-05 | SouthernBiotech | https://resources.southernbiotech.com/techbul/1071.pdf |
| Goat anti-mouse IgG2c-HRP | RRID:AB_2794462  Cat:1078-05 | SouthernBiotech | https://resources.southernbiotech.com/techbul/1078.pdf |
| Q-plex Mouse Cytokine – Screen (16-plex) | Cat:110949MS | Quansys Biosciences, West Logan, UT, USA | https://www.quansysbio.com/wp-content/uploads/2022/01/Mouse-Cytokine-Screen-HS-16-Plex-01-22-lr.pdf |
| GolgiPlug | RRID:AB_2869014  Cat: 555029 | BD Science, San Jose, CA, USA | https://www.bdbiosciences.com/content/bdb/paths/generate-tds-document.au.555029.pdf |
| phorbol 12-myristate 13-acetate | Cat:356150010 | Thermofisher | https://assets.thermofisher.com/chem-specs-pdf/retrievePdf?rootSku=35615&sku=356150010 |
| Ionomycin | Cat: J62448.MCR | Thermofisher | https://assets.thermofisher.com/chem-specs-pdf/retrievePdf?rootSku=J62448&sku=J62448.MCR |
| Fixable viability dye eFluor 780 | Cat:65-0865-14 | Thermofisher | https://www.thermofisher.com/document-connect/document-connect.html?url=https%3A%2F%2Fassets.thermofisher.com%2FTFS-Assets%2FLSG%2Fmanuals%2F65-0865.pdf |
| Fc block | RRID:AB_394656  Cat:553142 | BD Science | https://www.bdbiosciences.com/content/bdb/paths/generate-tds-document.ca.553142.pdf |
| FOXP3 Monoclonal Antibody (FJK-16s), PE-Cyanine7, eBioscience™ | RRID: AB_891552  Cat: 25-5773-82 | Thermofisher | https://www.thermofisher.com/order/genome-database/dataSheetPdf?producttype=antibody&productsubtype=antibody_primary&productId=25-5773-82&version=251 |
| BUV395 Hamster Anti-Mouse CD3e | RRID:  AB_2738278  Cat: 563565 | BD Biosciences | https://www.bdbiosciences.com/content/bdb/paths/generate-tds-document.us.563565.pdf |
| BV510 Rat Anti-Mouse CD8b | RRID: AB_2739908  Cat: 740155 | BD Biosciences | https://www.bdbiosciences.com/content/bdb/paths/generate-tds-document.us.740155.pdf |
| BUV737 Rat Anti-Mouse IFN-γ | RRID: AB_2870098  Cat: 612769 | BD Biosciences | https://www.bdbiosciences.com/content/bdb/paths/generate-tds-document.us.612769.pdf |
| Alexa Fluor® 700 anti-mouse CD4 Antibody | RRID: AB_493701  Cat: 100536 | BioLegend | https://www.biolegend.com/en-us/products/alexa-fluor-700-anti-mouse-cd4-antibody-3386?pdf=true&displayInline=true&leftRightMargin=15&topBottomMargin=15&filename=Alexa%20Fluor%C2%AE%20700%20anti-mouse%20CD4%20Antibody.pdf&v=20220118043120 |
| APC anti-mouse IL-17A Antibody | RRID: AB_536018  Cat: 506916 | BioLegend | https://www.biolegend.com/en-us/products/apc-anti-mouse-il-17a-antibody-3540?pdf=true&displayInline=true&leftRightMargin=15&topBottomMargin=15&filename=APC%20anti-mouse%20IL-17A%20Antibody.pdf&v=20220831123135 |
| Brilliant Violet 421™ anti-mouse IL-4 Antibody | RRID: AB_2562594  Cat: 504127 | BioLegend | https://www.biolegend.com/en-us/products/brilliant-violet-421-anti-mouse-il-4-antibody-7306?pdf=true&displayInline=true&leftRightMargin=15&topBottomMargin=15&filename=Brilliant%20Violet%20421%E2%84%A2%20anti-mouse%20IL-4%20Antibody.pdf&v=20220418063134 |
| FITC anti-mouse CD69 Antibody | RRID: AB_313108  Cat: 104505 | BioLegend | https://www.biolegend.com/en-us/products/fitc-anti-mouse-cd69-antibody-264?pdf=true&displayInline=true&leftRightMargin=15&topBottomMargin=15&filename=FITC%20anti-mouse%20CD69%20Antibody.pdf&v=20211217095219 |
| PerCP/Cyanine5.5 anti-mouse TNF-α Antibody | RRID: AB_961434  Cat: 506322 | BioLegend | https://www.biolegend.com/en-us/products/percp-cyanine5-5-anti-mouse-tnf-alpha-antibody-4438?pdf=true&displayInline=true&leftRightMargin=15&topBottomMargin=15&filename=PerCP/Cyanine5.5%20anti-mouse%20TNF-%CE%B1%20Antibody.pdf&v=20220606094049 |
| PE anti-human/mouse Granzyme B Recombinant Antibody | RRID: AB_2687032  Cat: 372208 | BioLegend | https://www.biolegend.com/en-us/products/pe-anti-human-mouse-granzyme-b-recombinant-antibody-14431?pdf=true&displayInline=true&leftRightMargin=15&topBottomMargin=15&filename=PE%20anti-human/mouse%20Granzyme%20B%20Recombinant%20Antibody.pdf&v=20220420084754 |
| Fixation Buffer | RRID: AB_2869005  Cat: 554655 | BD Science | https://www.bdbiosciences.com/content/bdb/paths/generate-tds-document.au.554655.pdf |
| Permeabilization buffer | RRID:  AB_2869011  Cat: 554723 | BD Science | https://www.bdbiosciences.com/content/bdb/paths/generate-tds-document.au.554723.pdf |
| PepTivator SARS-CoV-2 Prot_S | Cat: 130-126-700 | Miltenyi Biotec, Bergisch Gladbach, North Rhine-Westphalia, Germany | <https://www.miltenyibiotec.com/CA-en/products/peptivator-sars-cov-2-prot-s-107090.html#130-126-700>  <https://www.miltenyibiotec.com/upload/assets/IM0024612.PDF> |
| PepTivator SARS-CoV-2 Prot_S Complete | Cat: 130-127-951 | Miltenyi Biotec | <https://www.miltenyibiotec.com/CA-en/products/peptivator-sars-cov-2-prot-s-complete.html#130-127-951>  <https://www.miltenyibiotec.com/upload/assets/IM0027281.PDF> |
| SARS-CoV-2 Spike (Wuhan) | Produced In-lab | National Research Council of Canada, Montreal, QC, Canada | Akache, B., Renner, T.M., Stuible, M. *et al.* Immunogenicity of SARS-CoV-2 spike antigens derived from Beta & Delta variants of concern. *npj Vaccines* **7**, 118 (2022). https://doi.org/10.1038/s41541-022-00540-7 |
| SARS-CoV-2 RBD (Wuhan) | Produced In-lab | National Research Council of Canada, Montreal, QC, Canada | Colwill K, Galipeau Y, Stuible M, Gervais C, Arnold C, Rathod B, Abe KT, Wang JH, Pasculescu A, Maltseva M, Rocheleau L, Pelchat M, Fazel-Zarandi M, Iskilova M, Barrios-Rodiles M, Bennett L, Yau K, Cholette F, Mesa C, Li AX, Paterson A, Hladunewich MA, Goodwin PJ, Wrana JL, Drews SJ, Mubareka S, McGeer AJ, Kim J, Langlois MA, Gingras AC, Durocher Y. A scalable serology solution for profiling humoral immune responses to SARS-CoV-2 infection and vaccination. Clin Transl Immunology. 2022 Mar 23;11(3):e1380. doi: 10.1002/cti2.1380. PMID: 35356067; PMCID: PMC8942165.  Forest-Nault, C., Koyuturk, I., Gaudreault, J. et al. Impact of the temperature on the interactions between common variants of the SARS-CoV-2 receptor binding domain and the human ACE2. Sci Rep 12, 11520 (2022). https://doi.org/10.1038/s41598-022-15215-5 |
| SARS-CoV-2 Spike (B.1.1.7) | Produced In-lab | National Research Council of Canada, Montreal, QC, Canada | Akache, B., Renner, T.M., Stuible, M. *et al.* Immunogenicity of SARS-CoV-2 spike antigens derived from Beta & Delta variants of concern. *npj Vaccines* **7**, 118 (2022). https://doi.org/10.1038/s41541-022-00540-7 |
| SARS-CoV-2 Spike (B.1.351) | Produced In-lab | National Research Council of Canada, Montreal, QC, Canada | Akache, B., Renner, T.M., Stuible, M. *et al.* Immunogenicity of SARS-CoV-2 spike antigens derived from Beta & Delta variants of concern. *npj Vaccines* **7**, 118 (2022). https://doi.org/10.1038/s41541-022-00540-7 |
| SARS-CoV-2 Spike (P.1) | Produced In-lab | National Research Council of Canada, Montreal, QC, Canada | Akache, B., Renner, T.M., Stuible, M. *et al.* Immunogenicity of SARS-CoV-2 spike antigens derived from Beta & Delta variants of concern. *npj Vaccines* **7**, 118 (2022). https://doi.org/10.1038/s41541-022-00540-7 |
| SARS-CoV-2 Spike (B.1.617.2) | Produced In-lab | National Research Council of Canada, Montreal, QC, Canada | Akache, B., Renner, T.M., Stuible, M. *et al.* Immunogenicity of SARS-CoV-2 spike antigens derived from Beta & Delta variants of concern. *npj Vaccines* **7**, 118 (2022). https://doi.org/10.1038/s41541-022-00540-7 |
| SARS-CoV-2 Spike (B.1.1.529) | Produced In-lab | National Research Council of Canada, Montreal, QC, Canada | Akache, B., Renner, T.M., Stuible, M. *et al.* Immunogenicity of SARS-CoV-2 spike antigens derived from Beta & Delta variants of concern. *npj Vaccines* **7**, 118 (2022). https://doi.org/10.1038/s41541-022-00540-7 |
| SARS-CoV-2 Spike (BA.2) | Cat: NR-56517 | BEI Resources, Manassas, VA, USA | https://www.beiresources.org/Catalog/BEIProteins/NR-56517.aspx |
| Rockland Immunochemicals Rabbit TrueBlot: Anti-Rabbit IgG HRP - 18-8816-31 | Cat:  18-8816-31 | Rockland Immunochemicals, Pottstown, PA, USA | https://www.rockland.com/categories/trueblot/rabbit-trueblot-anti-rabbit-igg-hrp-18-8816-31/GetProductDataSheet/?code=18-8816-31 |
| BCG Danish | Cat: 35733 | ATCC | https://www.atcc.org/api/pdf/product-sheet?id=35733 |

**Supplemental Table 3. Table with Ad(Spike) primers**

| Primer Name | Primer Sequence |
| --- | --- |
| Spike_21_F | AGATCTGCCACCATGTTCGT |
| Spike_444_F | GATCCCTTCCTGGGTGTGTA |
| Spike_949_F | TCACTGTCGAGAAGGGCATA |
| Spike_1443_F | CAGAGATTTACCAAGCCGGTAG |
| Spike_1946_F | CCGGTAGCAATGTATTCCAGA |
| Spike_2453_F | CGACCCTTCCAAACCTTCTA |
| Spike_2943_F | GGCGCTATTTCCAGTGTTCT |
| Spike_3437_F | TACCGTGTACGATCCCCTTC |
| Spike_284_R | AAGGACGGGATTATCGAACC |
| Spike_792_R | AGTCGCCTGGTGTGAGGTAG |
| Spike_1290_R | TGCTATTTTCCCAGTTTGTCC |
| Spike_1803_R | TGCATGGGGTGATATCCAGT |
| Spike_2274_R | CGGTACTGTCACCGCAAATA |
| Spike_2804_R | GCAATCAGTTTCTGGTTCTCG |
| Spike_3297_R | AAGTGTGCCTTCCCATCGT |
| Spike_3784_R | CCACAGGAACAACACCCTTT |
| Spike_3954_R | CCCATATGTCCTTCCGAGTG |

**Supplemental Table 4. Serum neutralizing antibodies reported according to manufacturer’s guidelines**

| PBS | BCG(e)+Ad(e) | Ad(Spike) | BCG(e)+Ad(Spike) |
| --- | --- | --- | --- |
| NOT DETECTED | NOT DETECTED | DETECTED, >185 U/mL | DETECTED, >185 U/mL |
| NOT DETECTED | NOT DETECTED | DETECTED, >185 U/mL | DETECTED, >185 U/mL |
| NOT DETECTED | NOT DETECTED | DETECTED, >185 U/mL | DETECTED, >185 U/mL |
| NOT DETECTED | NOT DETECTED | DETECTED, >185 U/mL | DETECTED, >185 U/mL |
| NOT DETECTED | NOT DETECTED | DETECTED, >185 U/mL | DETECTED, >185 U/mL |
| NOT DETECTED | NOT DETECTED | DETECTED, >185 U/mL | DETECTED, >185 U/mL |
| NOT DETECTED |  | DETECTED, >185 U/mL | DETECTED, >185 U/mL |

**Supplemental Table 5. BALF neutralizing antibodies reported according to manufacturer’s guidelines**

| PBS | BCG(e)+Ad(e) | Ad(Spike) | BCG(e)+Ad(Spike) |
| --- | --- | --- | --- |
| NOT DETECTED | NOT DETECTED | DETECTED, 125.0 U/mL | DETECTED, 112.2 U/mL |
| NOT DETECTED | NOT DETECTED | DETECTED, >185 U/mL | DETECTED, >185 U/mL |
| NOT DETECTED | NOT DETECTED | DETECTED, >185 U/mL | DETECTED, >185 U/mL |
| NOT DETECTED | NOT DETECTED | DETECTED, >185 U/mL | DETECTED, >185 U/mL |
| NOT DETECTED | NOT DETECTED | DETECTED, <47 U/mL | DETECTED, 155.1 U/mL |
| NOT DETECTED | NOT DETECTED | DETECTED, >185 U/mL | DETECTED, >185 U/mL |
| NOT DETECTED |  | DETECTED, >185 U/mL | DETECTED, >185 U/mL |
