## Supplementary material for "BCG administration promotes the long-term protection afforded by a single-dose intranasal adenovirus-based SARS-CoV-2 vaccine": MDAR Checklist

### Materials Design Analysis Reporting (MDAR)
### Checklist for Authors

The MDAR framework establishes a minimum set of requirements in transparent reporting applicable to studies in the life sciences (see Statement of Task: doi:10.31222/osf.io/9sm4x.). The MDAR checklist is a tool for authors, editors, and others seeking to adopt the MDAR framework for transparent reporting in manuscripts and other outputs. Please refer to the MDAR Elaboration Document for additional context for the MDAR framework.

#### For all that apply, please note where in the manuscript the required information is provided.

#### Materials:

| Newly created materials | indicate where provided: page no/section/legend) | n/a |
|---|---|---|
| The manuscript includes a dedicated "materials availability statement" providing transparent disclosure about availability of newly created materials including details on how materials can be accessed and describing any restrictions on access. | At the end of the manuscript in the section Data and materials availability | |

| Antibodies | indicate where provided: page no/section/legend) | n/a |
|---|---|---|
| For commercial reagents, provide supplier name, catalogue number and RRID, if available. | In the Materials and Methods and supplementary materials | |

| DNA and RNA sequences | indicate where provided: page no/section/legend) | n/a |
|---|---|---|
| **Short novel DNA or RNA including primers, probes:** Sequences should be included or deposited in a public repository. | In the supplementary materials | |

| Cell materials | indicate where provided: page no/section/legend | n/a |
|---|---|---|
| **Cell lines:** Provide species information, strain. Provide accession number in repository **OR** supplier name, catalog number, clone number, **OR** RRID. | In the Materials and Methods and supplementary materials | |
| **Primary cultures:** Provide species, strain, sex of origin, genetic modification status. | In the Materials and Methods and supplementary materials | |

| Experimental animals | indicate where provided: page no/section/legend) | n/a |
|---|---|---|
| **Laboratory animals or Model organisms:** Provide species, strain, sex, age, genetic modification status. Provide accession number in repository **OR** supplier name, catalog number, clone number, **OR** RRID. | In the Materials and Methods | |
| **Animal observed in or captured from the field:** Provide species, sex, and age where possible. | | n/a |

| Plants and microbes | indicate where provided: page no/section/legend) | n/a |
|---|---|---|
| **Plants:** provide species and strain, ecotype and cultivar where relevant, unique accession number if available, and source (including location for collected wild specimens). | | n/a |
| **Microbes:** provide species and strain, unique accession number if available, and source. | In the Materials and Methods and supplementary materials | |

| Human research participants | indicate where provided: page no/section/legend) or state if these demographics were not collected | n/a |
|---|---|---|
| If collected and within the bounds of privacy constraints report on age, sex and gender or ethnicity for all study participants. | | n/a |

#### Design:

| **Study protocol** | **indicate where provided: page no/section/legend)** | **n/a** |
|---|---|---|
| If study protocol has been pre-registered, provide DOI. For clinical trials, provide the trial registration number **OR** cite DOI. | | n/a |

| **Laboratory protocol** | **indicate where provided: page no/section/legend)** | **n/a** |
|---|---|---|
| Provide DOI **OR** other citation details if detailed step-by-step protocols are available. | | n/a |

| **Experimental study design (statistics details)** | | |
|---|---|---|
| **For in vivo studies:** State whether and how the following have been done | **indicate where provided: page no/section/legend. If it could have been done, but was not, write not done** | **n/a** |
| Sample size determination | In the Materials and Methods within Study design | |
| Randomisation | In the Materials and Methods within Study design | |
| Blinding | In the Materials and Methods within Study design | |
| Inclusion/exclusion criteria | In the Materials and Methods within Study design | |

| **Sample definition and in-laboratory replication** | **indicate where provided: page no/section/legend** | **n/a** |
|---|---|---|
| State number of times the experiment was replicated in laboratory. | In Materials and Methods | |
| Define whether data describe technical or biological replicates. | In all relevant figure captions | |

| **Ethics** | **indicate where provided: page no/section/legend** | **n/a** |
|---|---|---|
| **Studies involving human participants:** State details of authority granting ethics approval (IRB or equivalent committee(s), provide reference number for approval. | | n/a |
| **Studies involving experimental animals:** State details of authority granting ethics approval (IRB or equivalent committee(s), provide reference number for approval. | In Materials and Methods within Animal ethics | |
| **Studies involving specimen and field samples:** State if relevant permits obtained, provide details of authority approving study; if none were required, explain why. | | n/a |

| **Dual Use Research of Concern (DURC)** | **indicate where provided: page no/section/legend** | **n/a** |
|---|---|---|
| If study is subject to dual use research of concern regulations, state the authority granting approval and reference number for the regulatory approval. | | n/a |

#### Analysis:

| Attrition | indicate where provided: page no/section/legend | n/a |
|---|---|---|
| Describe whether exclusion criteria were preestablished. Report if sample or data points were omitted from analysis. If yes report if this was due to attrition or intentional exclusion and provide justification. | In the Materials and Methods within Study design | |

| Statistics | indicate where provided: page no/section/legend | n/a |
|---|---|---|
| Describe statistical tests used and justify choice of tests. | In the Materials and Methods within Statistical analysis | |

| Data availability | indicate where provided: page no/section/legend | n/a |
|---|---|---|
| For newly created and reused datasets, the manuscript includes a data availability statement that provides details for access or notes restrictions on access. | At the end of the manuscript in the section Data and materials availability | |
| If newly created datasets are publicly available, provide accession number in repository **OR** DOI **OR** URL and licensing details where available. | | n/a |
| If reused data is publicly available provide accession number in repository **OR** DOI **OR** URL, **OR** citation. | | n/a |

| Code availability | indicate where provided: page no/section/legend | n/a |
|---|---|---|
| For all newly generated custom computer code/software/mathematical algorithm or re-used code essential for replicating the main findings of the study, the manuscript includes a data availability statement that provides details for access or notes restrictions. | | n/a |
| If newly generated code is publicly available, provide accession number in repository, **OR** DOI **OR** URL and licensing details where available. State any restrictions on code availability or accessibility. | | n/a |
| If reused code is publicly available provide accession number in repository **OR** DOI **OR** URL, **OR** citation. | | n/a |

#### Reporting

MDAR framework recommends adoption of discipline-specific guidelines, established and endorsed through community initiatives. Journals have their own policy about requiring specific guidelines and recommendations to complement MDAR.

| **Adherence to community standards** | **indicate where provided: page no/section/legend** | **n/a** |
|---|---|---|
| State if relevant guidelines (e.g., ICMJE, MIBBI, ARRIVE) have been followed, and whether a checklist (e.g., CONSORT, PRISMA, ARRIVE) is provided with the manuscript. | ARRIVE (Supplemental Data) | |
